# Principles of visual cortex excitatory microcircuit organization

**DOI:** 10.1101/2023.12.30.573666

**Authors:** Christina Y.C. Chou, Hovy H.W. Wong, Connie Guo, Kiminou E. Boukoulou, Cleo Huang, Javid Jannat, Tal Klimenko, Vivian Y. Li, Tasha A. Liang, Vivian C. Wu, P. Jesper Sjöström

**Affiliations:** Centre for Research in Neuroscience, Brain Repair and Integrative Neuroscience Program, Department of Neurology and Neurosurgery, The Research Institute of the McGill University Health Centre, Montreal, Quebec, H3G 1A4, Canada; Integrated Program in Neuroscience, McGill University, Montreal, Quebec, H3A 2B4, Canada

**Keywords:** Connectivity, Short-term plasticity, Optogenetics, Microcircuit, Plasticitome

## Abstract

Microcircuit function is determined by patterns of connectivity and short-term plasticity that vary with synapse type. Elucidating microcircuit function therefore requires synapse-specific investigation. The state of the art for synapse-specific measurements has long been paired recordings. Although powerful, this method is slow, leading to a throughput problem. To improve yield, we therefore created optomapping — an approximately 100-fold faster 2-photon optogenetic method — which we validated with paired-recording data. Using optomapping, we tested 15,433 candidate excitatory inputs to find 1,184 connections onto pyramidal, basket, and Martinotti cells in mouse primary visual cortex, V1. We measured connectivity, synaptic weight, and short-term dynamics across the V1 layers. We found log-normal synaptic strength distributions, even in individual inhibitory cells, which was previously not known. We reproduced the canonical circuit for pyramidal cells but found surprising and differential microcircuit structures, with excitation of basket cells concentrated to layer 5, and excitation of Martinotti cells dominating in layer 2/3. Excitation of inhibitory cells was denser, stronger, and farther-reaching than excitation of excitatory cells, which promotes stability and difference-of-Gaussian connectivity. We gathered an excitatory short-term plasticitome, which revealed that short-term plasticity is simultaneously target-cell specific and dependent on presynaptic cortical layer. Peak depolarization latency in pyramidal cells also emerged as more heterogeneous, suggesting heightened sensitivity to redistribution of synaptic efficacy. Optomapping additionally revealed high-order connectivity patterns including shared-input surplus for interconnected pyramidal cells in layer 6. Optomapping thus offered both resolution to the throughput problem and novel insights into the principles of neocortical excitatory fine structure.

**HIGHLIGHTS:**

- 2-photon optomapping of microcircuits is verified as fast, accurate, and reliable
- Synaptic weights distribute log-normally even for individual inhibitory neurons
- Maximal excitation of basket and Martinotti cells in layer 5 and 2/3, respectively
- Short-term plasticity depends on layer in addition to target cell

## INTRODUCTION

Information processing in the brain is determined by the patterns of connectivity as well as the short-term plasticity at synapses among different types of neurons.^1-3^ Connectivity patterns are established by a combination of nature and nurture,^4^ i.e., genetic information and experience-dependent synaptic plasticity.^5-7^

In the textbook view, canonical cortical microcircuits are organized in the form of columns and pathways between different cortical layers.^8-10^ Thalamic pathways chiefly innervate layer 4 (L4), the input layer. Ascending projections from L4 largely terminate in L2/3, the computational layer.^10,11^ L2/3 PCs then project to L5, and L5 PCs project to L6 or subcortical regions.^10,11^ Simplistically, L5 is thus considered the principal output layer.^8,9,11,12^ However, because many classic studies explored synaptic pathways rather than individual connections, there is a relative paucity of information on connectivity and short-term plasticity with synapse-type specificity.^13-15^

Moreover, the rules that govern plasticity and connectivity patterns are specific to synapse type.^13-15^ For instance, the same neocortical pyramidal cell (PC) produces short-term depressing or facilitating synaptic outputs depending on target cell type.^16,17^ Such synapse-type-specific patterning is expected, since different neuronal classes play drastically different functional roles, e.g., mediating excitation or inhibition.^18-20^

Cortical microcircuits are furthermore populated by recurring connectivity motifs established with key inhibitory interneuron types. For instance, basket cells (BCs)^18-20^ mediate fast-onset perisomatic inhibition of PCs,^21^ which enhances the temporal fidelity of PC spiking.^22^ Martinotti cells (MCs),^18-20^ on the other hand, provide slow-onset inhibition of PC apical dendrites,^21,23^ which synchronizes spiking of PC populations by negative feedback.^24^ Interneurons such as BCs and MCs can thereby dynamically reroute information flow across the PC somato-dendritic axis.^25^ Cortical PC computations are therefore critically determined by neighboring BCs and MCs.

It is thus important that cortical connections are explored with synapse-type specificity.^13-15^ To achieve such fine-grain resolution, multiple patch clamp has long remained the state-of-the-art method for recording connected neuronal pairs. In the mouse, this powerful approach has, e.g., revealed the non-random structure of the L5 PC network,^26^ the functional specificity of the L2/3 PC network,^27,28^ the excitatory synaptic organization of developing barrel cortex,^29^ and the excitatory as well as inhibitory adult V1 circuits,^30,31^ even with direct comparison to the human counterpart.^30^

Multiple patch, however, suffers from sampling shortcomings. For instance, results are gathered across cells, acute slices, and animals, typically over months and years.^26,27,29-31^ It thus remains possible that findings arise as a consequence of pooling data across experiment days. For example, high-order connectivity patterns have often been found by sampling across paired recordings over many experimental days.^26,32,33^ Similarly, it is not clear if long-tailed log-normal distributions of synaptic weights^26,34^ holds true for individual cortical neurons or if it emerges as an artifact of cross-cell sampling, which is important as log-normality underlies functions such as feature preference.^27,28^

Furthermore, because multiple patch is prohibitively slow, it is generally not feasible to measure connectivity or plasticity across the entire thickness of a given cortical area, which has limited what kinds of queries neuroscientists can explore. There has thus been a long-standing need for approaches with considerably higher throughput.^13,35^

To resolve this throughput problem, we created and validated optomapping, a high-throughput circuit charting approach based on 2-photon (2P) optogenetics combined with patch-clamp electrophysiology. We optomapped connectivity, synaptic weights, and short-term dynamics of excitatory synapses onto PCs, BCs, and MCs across the layers of mouse V1, which revealed striking and surprising target-cell-specific differences. Our findings provide fresh perspective on the principles that govern V1 excitatory fine structure.

## RESULTS

### Optogenetic activation of candidate presynaptic neurons

To link postsynaptic responses to individual presynaptic cells, we sought to reliably photoactivate spiking in candidate presynaptic neurons with single-cell resolution and millisecond temporal precision. To optogenetically stimulate neurons with high spatial resolution, researchers have successfully turned to 2P excitation, because it limits photoexcitation to a diffraction-limited femtoliter volume.^36^

To drive spiking with 2P excitation, we expressed a soma-targeted version of the red-shifted high-current opsin ChroME^37^ in neocortical PCs by viral infection with AAV-CAG-DIO-ChroME-ST-P2A-H2B-mRuby3 in Emx1^Cre/Cre^ mice.^38^ To discriminate interneurons from PCs, we also tagged interneurons by co-infecting with AAV-mDlx-GFP-Fishell-1.^39^ These viruses were injected into V1 on postnatal day (P) 0-2,^40^ which gave rise to robust, non-overlapping mRuby and GFP expression by P18 and onwards (Figures 1A and 1B).

**Figure 1.**
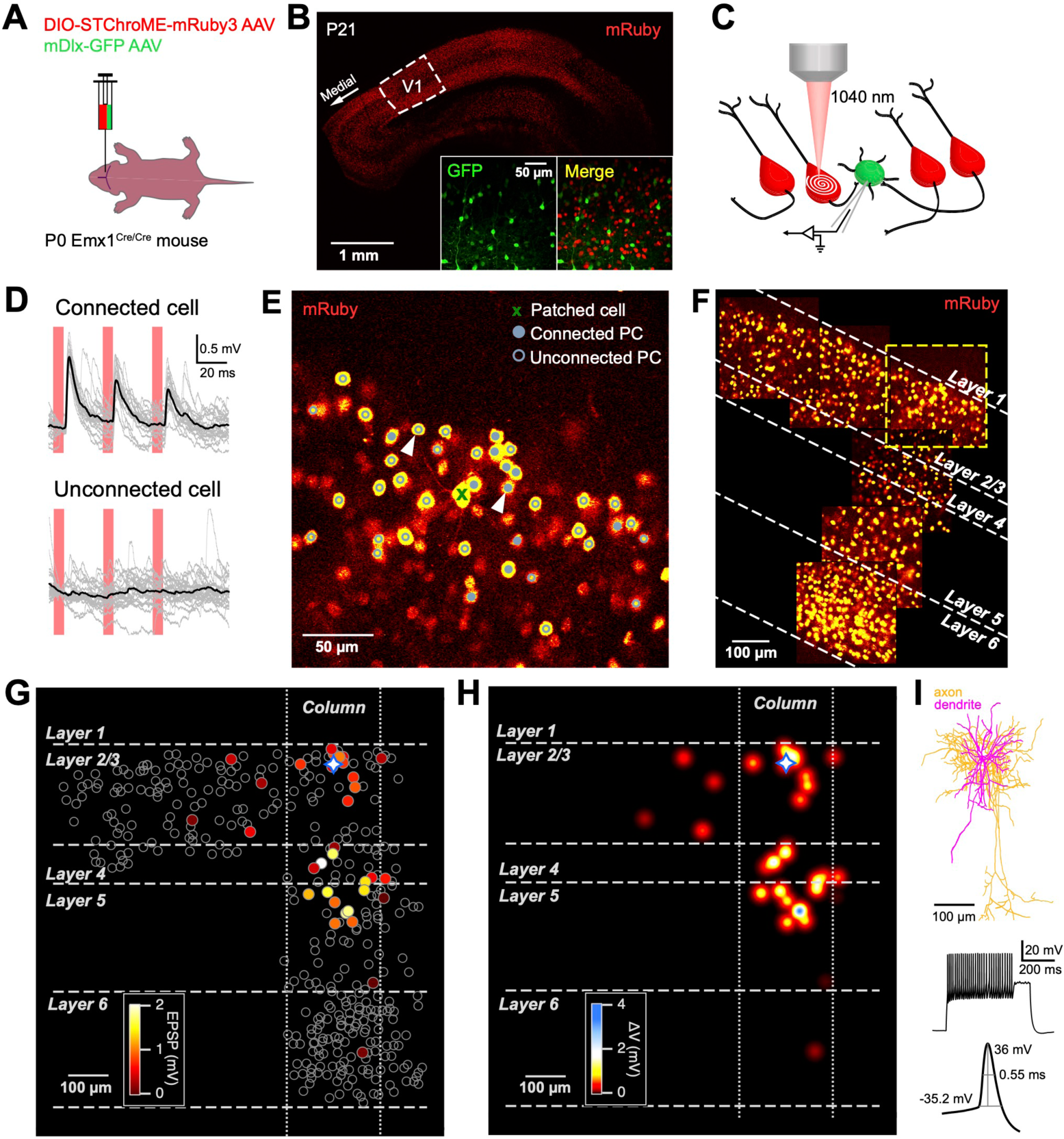
Sample optomap of excitatory inputs onto an individual cell. (A) Using neonatal viral injection^40^ in V1 of Emx1^Cre/Cre^ mice^38^, we expressed the soma-targeted ChroME opsin with mRuby label^37^ in PCs as well as GFP in interneurons.^39^ (B) In this acute slice cut at P21, nonoverlapping mRuby and GFP expression were observed, consistent with specific labelling of PCs and interneurons. (C) To find connected PCs, we spiral scanned mRuby-positive cells with a 1040-nm 2P laser beam while whole-cell recording a postsynaptic cell, here a GFP-expressing interneuron. (D) Sample responses from connected and unconnected PCs. Based on statistically significant depolarization across 20 individual traces (grey), averaged traces (black) were semi-automatically deemed to contain EPSPs or not (see Methods). (E) In this sample FOV, spiral scanning mRuby-positive cells either elicited EPSPs (closed circles) or not (open circles) in the patch-clamped postsynaptic cell in L2/3 (green X). Arrowheads correspond to sample traces in (D). Spiral scans were semi-automatically positioned on mRuby-positive cells by software algorithm (see Methods). (F) By repeating this process for neighbouring FOVs, we optomapped the entire cortical thickness as well as hundreds of µm laterally from the patched postsynaptic cell. Yellow dash: FOV in (E). White dash: layer boundaries. (G) We colour-coded the location of presynaptic PCs by the EPSP amplitude (circles) elicited in the patched L2/3 cell (diamond). To enable fair cross-layer comparison for different postsynaptic cells, a 200-µm-wide column (vertical dotted lines) was centred on the patched cell. (H) For the sample in (G), we generated a synaptic input density map for the patched L2/3 cell (diamond). Density maps enabled averaging across different postsynaptic cells of the same class (see main text). (I) Based on morphology and spike pattern, the postsynaptic cell was classified as an inhibitory BC (for cell classification approach, see main text and Methods).

We first made sure that we could reliably drive spiking of ChroME-expressing PCs with 2P excitation. To elicit strong opsin currents, we covered the target soma with light by spiral-scanning it with a 1040-nm 2P beam for 7 ms (Figure S1A), which evoked spiking with 100% fidelity up to ∼70 Hz (Figures S1E and S1F). The probability of eliciting surplus spikes was indistinguishable from that with 5-ms-long current pulses (Figures S1G and S1H). The spike latency was ∼5 ms and the jitter was below 0.5 ms (Figures S1B-S1D). We could thus reliably drive PC spiking with high temporal resolution.

Although 2P microscopy lateral resolution is high (typically <0.5 µm), the axial resolution can be appreciably worse.^36^ This is in particular a concern for 2P photostimulation, since we aimed to activate only one presynaptic candidate cell at a time. We therefore assessed the axial resolution by offsetting in z while spiral-scanning centered on a patched PC soma. We found that the probability distribution of light-evoked spiking at half-maximum had a half-width *w*_½_ of ∼10 µm (Figure S1I), which corresponds well to the ∼10-µm radius of a PC soma in mouse neocortex,^41^ as well as the ∼20 µm intercell distance of neurons in the cortex (see Methods).

Taken together, our findings demonstrate that by spiral-scanning a femtosecond laser beam over ChroME-expressing neurons, we could reliably drive PC spiking with single-cell resolution and millisecond temporal precision (Figure S1). These results agree with previous reports on 2P optogenetics.^37,42-45^

### Sample optogenetic map of synaptic connections

To map connectivity within a 265×265-µm FOV, we patched a postsynaptic PC, BC, or MC and subsequently activated candidate presynaptic neurons by sequentially spiral-scanning mRuby3-positive cells in a single focal plane (see sample BC in Figure 1C-1E). To measure short-term plasticity, each mRuby3-positive cell was subjected to a train of three spiral scans at 30 Hz. To assess response variability and coefficient of variation (CV), the sequential optogenetic stimulation of mRuby3-positive cells in a FOV was repeated 20 times every 20-40 seconds. To avoid artifacts related to acute slice preparation, we carried out optomapping ∼100 µm from the slice surface (Figure S2; see Methods). To optomap the entire cortical thickness as well as hundreds of µm laterally to the recorded cell, this process was repeated for nearby FOVs while continuously recording the postsynaptic cell (Figure 1F). Typically, this process took 5-15 min per FOV depending on the number of mRuby3-positive cells (total time for Figure 1F: 51 min; on average: 72 ± 10 min, n = 6 randomly selected experiments).

In offline analysis, presynaptic PCs that consistently elicited EPSPs in the patched cell across the 20 trials were semi-automatically tagged as connected (Figure 1C; see Methods). EPSP amplitude was used as a metric of connective strength and the paired-pulse ratio (PPR) as a quantification of short-term plasticity (see below and Methods). To enable comparison of connectivity and synaptic strength across cortical layers, we assigned a 200-µm-wide vertical column centered on the postsynaptic cell as well as layer boundaries based on standard layer-specific features (see Methods).^46-48^

In this sample experiment, the high throughput of optomapping allowed us to probe 363 candidate presynaptic PCs, of which 35 were connected (Figure 1G). For comparison, this throughput over a whole-cell recording lasting 1-2 hours is ∼100-fold faster than multiple patch-clamp recordings, depending on specific comparison.^26,27,29-31^ To enable averaging of connectivity maps across different postsynaptic cells of the same type, we created synaptic input density maps (Figure 1H; see Methods). For context, such maps cannot be meaningfully created for paired recordings.

From this individual map, we observed that, although this L2/3 BC received appreciable within-layer excitation as previously reported,^30,49^ cross-layer excitation from L4 and L5 was substantial, revealing high success rate at finding connections hundreds of µm from the patched cell. Laterally in L2/3, connectivity qualitatively died down over tens to hundreds of µm (Figures 1G and 1H).^30,49^ Detailed statistical comparisons across different postsynaptic cells of these observations are provided below.

### Accounting for optogenetic stimulation artifacts

Emx1^Cre/Cre^ mice drive expression in >90% of excitatory neurons.^38,50^ Consequently, when the postsynaptic cell was a PC, we occasionally directly depolarized it when spiral-scanning nearby presynaptic PCs. However, compared to EPSPs, direct depolarization was instantaneous, had small CV, and no short-term depression (Figure S3). Furthermore, direct depolarization only occurred within ∼60 µm of the patched PC (Figure S3E-H). We thus relied on a combination of patched cell type (PC vs. BC/MC), depolarization onset latency, stimulation location, and the analysis of CV and short-term plasticity to determine whether a stimulation location elicited direct depolarization (Figure S3B-I), which we then accounted for (Figure S3J; Methods). Direct activation affected <2% of input cells within a FOV (9 out of 493 photo-stimulation locations, n = 7 PCs), indicating a negligible impact.

In conclusion, we could reliably measure PC→PC connectivity even for nearby PCs. However, for a few inputs close to postsynaptic PCs, synaptic weight and dynamics might be distorted. Since Emx1-Cre does not target inhibitory cells,^38,50^ this artifact did not affect PC→BC or PC→MC optomapping.

### Identification of PCs, BCs, and MCs

Because PC axons target thousands of postsynaptic cells^8^ with differing functional properties,^18-20^ connectivity rates, synaptic weights, and short-term plasticity are expected to depend on the target cell type.^13-15^ We therefore independently categorized postsynaptic cells into PC, BC, and MC classes using hierarchical clustering based on morphological and electrophysiological characteristics (Figure S4).^51^ These characteristics were obtained by patch clamp followed by biocytin histology and confocal imaging (Figure 1I and S5; see Methods). Excitatory and inhibitory neuron properties were consistent with prior descriptions^18-20^ indicating that we accurately classified PCs, BCs, and MCs.

### Excitatory synaptic strengths distribute log-normally

Previous paired-recording studies demonstrated that synaptic strengths distributed log-normally,^26,28,29,52^ which is important as this may stimulus-specifically amplify functional responses.^27,28^ However, these studies pooled data across different cells and days.^26,28,29,52^ We therefore relied on optomapping to ask if synaptic weights onto individual neurons distribute log-normally. It has additionally been argued that log-normal weight distributions arise from Hebbian learning.^53^ However, PC→BC connections are known for anti-Hebbian plasticity,^54,55^ so we speculated that excitatory inputs onto BCs might not distribute log-normally. We tested this idea by assessing EPSP amplitude distributions in individual PCs, BCs, and MCs in L2/3 (Figure 2A).

**Figure 2.**
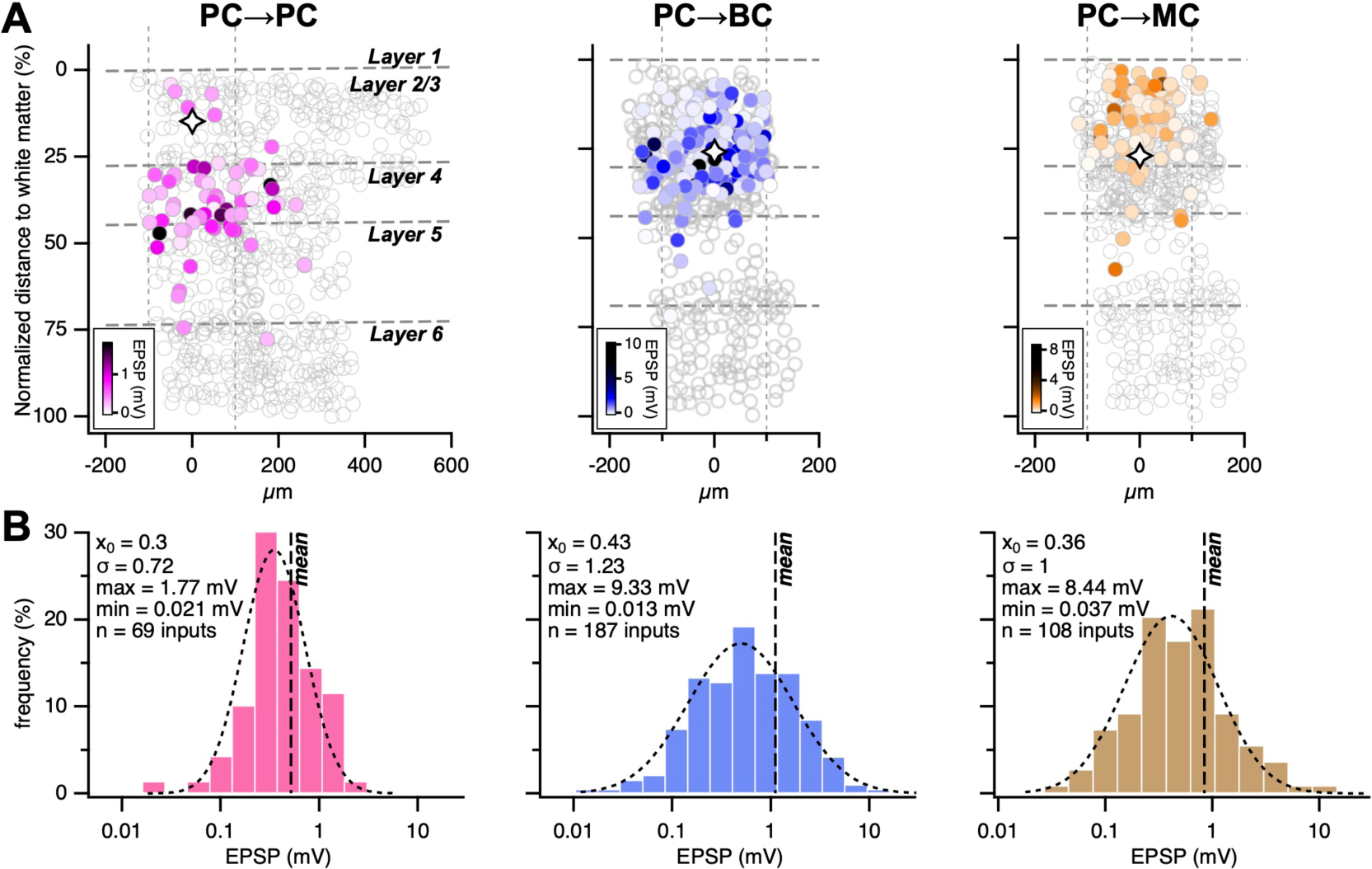
EPSPs in individual L2/3 cells distribute log-normally. (A) We optomapped dozens of excitatory inputs onto individual cells, which enabled us to explore the distribution of synaptic weights in a single L2/3 PC (left), BC (middle), and MC (right). Open circles: unconnected presynaptic neurons. Closed circles: connected presynaptic neurons. Diamond: patched postsynaptic cell. (B) EPSP distributions onto individual cells shown in (A) were best fit by a log-normal model (see Methods), as were the corresponding distributions pooled across cells of the same type (Figure S6). Dashed line: mean EPSP amplitude. Dotted line: log-normal fit.

As we previously described for L5 PCs,^26^ we observed many more weak connections than strong ones in L2/3. By comparing with different distribution types (see Methods), we found that the best fit was log-normal for individual PCs, BCs, as well as MCs (Figure 2B).

In agreement with prior studies,^26,29,52^ ensemble EPSP amplitudes pooled across PCs were also best fit by a log-normal distribution (Figure S6). For L6 PCs, best fits were sinh-arcsinh distribution, followed by Johnson’s *S_U_*, with log-normal in third place (Figure S6). However, these are all similar log-normal variants.

Excitatory inputs onto BCs and MCs in L2/3, L5, and L6 were also log-normally distributed (Figure S6). Although means and standard deviations differed across PC, BC, and MC types (see below), the best-fit distribution model did not.

From these findings emerged the principle that excitatory synaptic weights generally distribute log-normally, regardless of target cell type. Notably, excitatory inputs onto individual BCs and MCs also obey this rule, suggesting that log-normality is not a direct consequence of Hebbian learning.^53^

### In L2/3, PCs and BCs receive strong ascending drive while that of MCs is local

We next wondered if excitatory input spatial distributions differed across PC, BC, and MC types. In the neocortex, PC→PC connectivity within and across layers have been well studied using paired recordings,^26,27,29,30,49,52,56-60^ but PC→BC and PC→MC synapses have not been as extensively investigated, especially not with respect to dependence on cortical layer.^30,31,61,62^ Additionally, the complexity of paired patch has made it difficult to systematically explore how synaptic connections distribute spatially.^49,63^ High throughput optical methods are better suited for spatial mapping but classically did not have single-cell resolution.^64-67^ We therefore optomapped the spatial input distributions for PCs, BCs, and MCs, starting in L2/3.

In L2/3, strikingly different connectivity patterns emerged for PCs, BCs, and MCs (Figure 3A), which we carefully analyzed statistically with generalized linear mixed-models (GLMM) and linear mixed-models (LMMs; see Methods). Consistent with the canonical cortical circuit,^8-11^ L2/3 PCs received more inputs from L4 than from other layers (Figures 3A and 3B), but L5 PC→L2/3 PC connectivity was also high, suggesting potent excitatory feedback from the cortical output layer, L5, as previously reported.^68^ L6 PC→L2/3 PC connectivity was low. Similar to L2/3 PCs, L2/3 BCs received many excitatory inputs from L2/3, L4, and L5, but few from L6 (Figures 3A and B).

**Figure 3.**
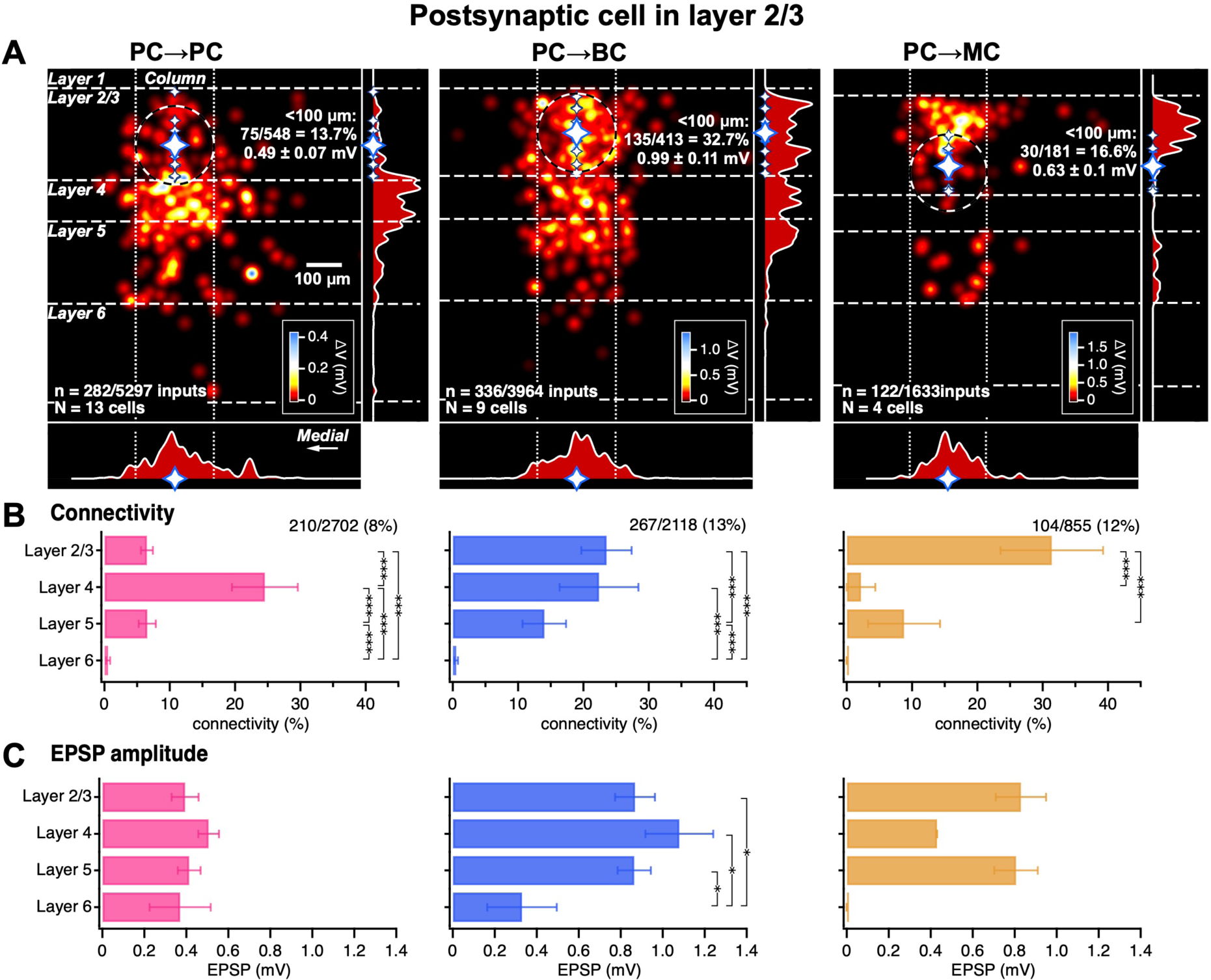
In L2/3, strong ascending PC→PC/BC drive but local PC→MC excitation. (A) Synaptic input density maps for L2/3 PCs (left), BCs (middle), and MCs (right) were generated by averaging individual maps (Figure 1H). Insets right and bottom show the vertical and horizontal density projections, respectively. Connectivity within a 100-µm radius (dashed circle) is shown to enable comparison with paired patch-clamp recordings, which typically are close.^49^ Dashed horizontal lines: layer boundaries. Dotted lines: vertical column. (B) L2/3 PCs had higher excitatory connectivity from L4 than from other layers, and lower from L6. L2/3 BCs had higher excitatory connectivity from L2/3 and L4. L2/3 MCs, on the other hand, had higher excitatory connectivity from L2/3 than from other layers, and with few L4 and L6 inputs.^70^ Comparing across cell types, L2/3 PCs were more frequently connected to L2/3 BCs and L2/3 MCs than to L2/3 PCs (PC vs. BC, p < 0.001, PC vs. MC, p < 0.001), L4 PCs were less frequently connected to L2/3 MCs than to L2/3 PCs and L2/3 BCs (PC vs. MC, p < 0.001, BC vs. MC, p < 0.001), and L5 PCs were more frequently connected to L2/3 BCs than to L2/3 PCs (PC vs. BC, p < 0.05), whereas for L6 PC inputs, we found no differences. To account for postsynaptic cell as a random variable in these comparisons, we used GLMM statistics (see Methods, presynaptic layer and cell type interaction effect, p < 0.001). (C) L2/3 PCs received weaker excitatory inputs than L2/3 BCs (PC vs. BC, p < 0.01), regardless of presynaptic layer. MCs, however, were indistinguishable in this regard (PC vs. MC, p = 0.15, BC vs. MC, p = 0.99). LMM statistics revealed that EPSP amplitudes depended on cell type (p < 0.01) but not on presynaptic layer (p = 0.12, interaction effect p = 0.74). Mean ± SEM \**P < 0.05*, \*\**P < 0.01*, \*\*\**P < 0.001*.

In contrast, L2/3 MCs received the most inputs from nearby PCs in the same layer, especially from L2 (Figure 3A), significantly more so than L2/3 PCs did. L2/3 MCs also received more inputs from L5 than from L4 (Figures 3A and 3B). In fact, L4 PC→L2/3 MC connectivity was lower than that of L4 PC→L2/3 BCs and L4 PC→L2/3 PCs.

L6→L2/3 excitatory connections were strikingly rare for all three cell types (Figure 3A and 3B). We did not observe a single L6 PC→L2/3 MC connection despite sampling 296 candidate L6 inputs (Figure 3B).

Laterally, connectivity died down over tens to hundreds of µm.^30,49^ There were few connectivity differences across the different cell types (Figure 3A), although excitation of BCs seemed to originate farther away (see below for detailed analysis). As noted in prior studies,^30,49^ excitatory inputs were chiefly detected within 300 µm lateral of the postsynaptic cell. Even though connections were sampled hundreds of microns laterally, only a few long-range connections were found (Figure 3A). We did not observe any mediolateral asymmetries (Figure 3A).

To compare synaptic strengths, we relied on the amplitude of the first EPSP in a train of three EPSPs (Figure 1D; see Methods). For L2/3 PCs and MCs, input strengths were indistinguishable across layers (Figure 3C). This was mostly true for L2/3 BCs as well, except synaptic weights from L6 were weaker (Figure 3C). Comparing across cell types, excitatory inputs were much stronger onto L2/3 BCs than onto L2/3 PCs and L2/3 MCs (Figures 3A and 3C). These comparisons, however, did not account for the well-known strong short-term facilitation of PC→MC connections,^16,17,46,69^ which means for PC→MC synapses, the last EPSP in a train of three (Figure 1D) was appreciably larger than the first EPSP. We revisit PC→MC short-term facilitation separately below.

Taken together, these findings reveal distinct spatial distributions of excitatory inputs onto PCs, BCs, and MCs in L2/3. For instance, we found strong ascending excitatory drive of PCs and BCs but mostly within-layer local excitation of MCs. Overall, excitation of BCs was also much more potent than that of PCs or MCs. L6 PCs were largely disconnected from these L2/3 neurons, suggesting computational independence.^30^

### In L5, most excitatory inputs originate from L5

We next explored the spatial distribution of cortex-wide excitatory inputs onto PCs, BCs, and MCs in L5. Here, BCs received more intralaminar inputs than L5 PCs and L5 MCs did (Figures 4A and 4B). Interestingly, L5 neurons of all three types received few excitatory inputs from L6, despite these layers being adjacent (Figures 4A and 4B). This echoes the finding that L6→L2/3 excitatory connections were rare (Figures 3A and 3B).

**Figure 4.**
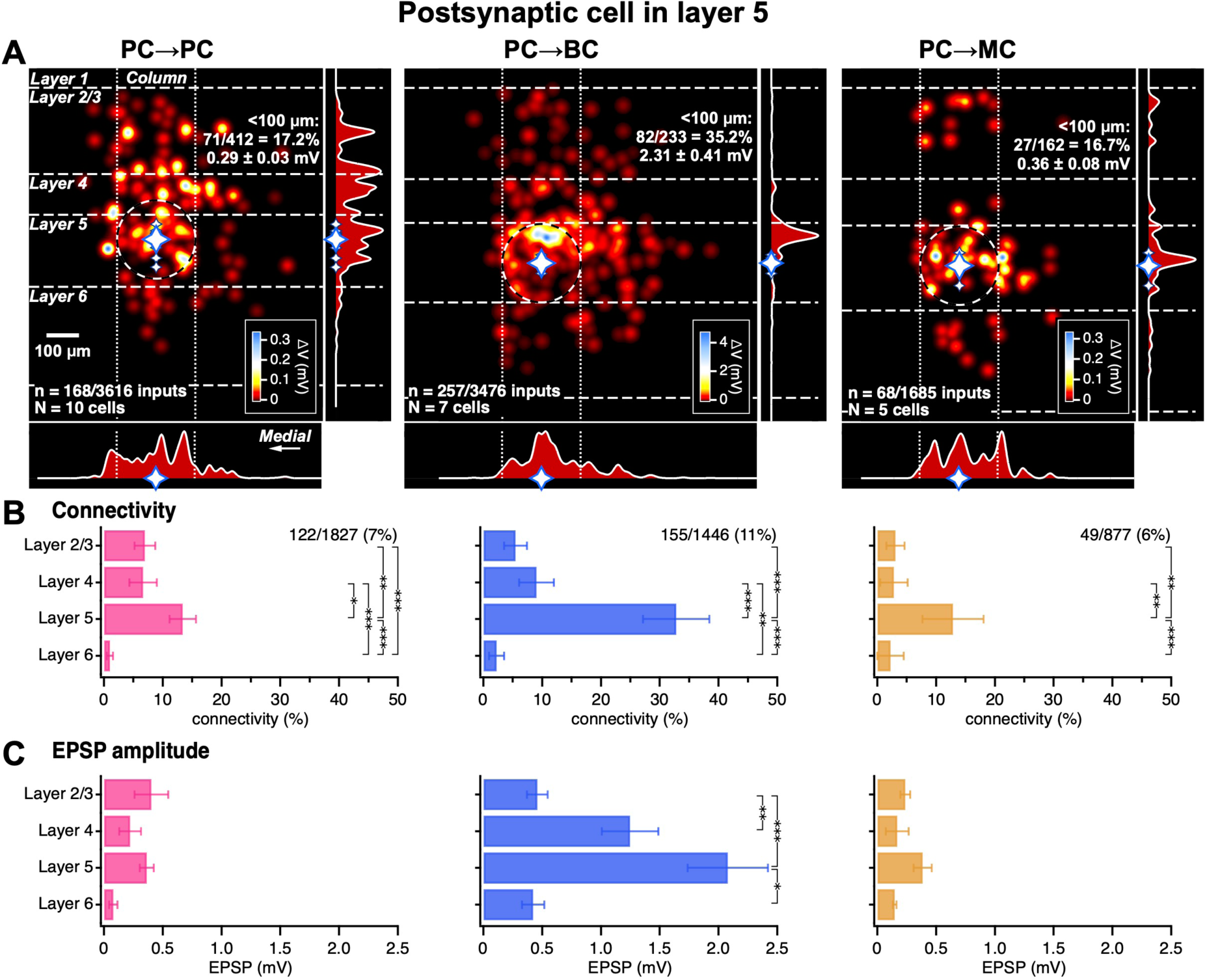
For PCs, BCs, and MCs in L5, excitatory drive concentrated to L5. (A) Synaptic input density maps for L5 PCs (left), BCs (middle), and MCs (right) were generated by averaging individual maps (Figure 1H). Insets right and bottom show the vertical and horizontal density projections, respectively. Connectivity within a 100-µm radius (dashed circle) simplifies comparison with paired patch-clamp recordings.^26,46^ Dashed horizontal lines: layer boundaries. Dotted lines: vertical column. (B) L5 PCs, BCs, and MCs all had higher excitatory connectivity from L5 than from L2/3, L4, or L6. Within L5, BCs received more excitatory inputs than PCs and MCs did (PCs vs. BCs, p < 0.001, BCs vs. MCs, p < 0.01). GLMM statistics revealed that connectivity depended on presynaptic layer and cell type (p < 0.001). (C) Excitatory synaptic input weights onto L5 PCs and L5 MCs did not differ by layer. L5 PC→L5 BC and L4 PC→L5 BC inputs were stronger than inputs from other layers. Inputs from L4 and L5 were stronger onto BCs than onto PCs and MCs (from L4, PCs vs. BCs, p < 0.05, BCs vs. MCs, p < 0.05; from L5, PCs vs. BCs, p < 0.01, BCs vs. MCs, p < 0.01). LMM statistics revealed that EPSP amplitudes depended on both cell type and on presynaptic layer (p < 0.01). Mean ± SEM. \**P < 0.05*, \*\**P < 0.01*, ** *< 0.001*.

In L5, connection strength depended on input layer for BCs, but not for PCs or MCs (Figure 4C), like in L2/3. Comparing across cell types, excitatory connections from L4 and L5 were stronger onto L5 BCs than onto L5 PCs or L5 MCs. Interestingly, PC→BC input weights concentrated to upper L5 (Figure 4A).

In sum, these findings reveal relatively similar spatial distributions of excitatory inputs onto PCs, BCs, and MCs in L5, with most excitation originating from within L5, and only some descending input. As for L2/3 (Figure 3), excitatory drive of BCs was stronger than that of PCs or MCs. L6 PC inputs were also relatively disconnected from all three L5 neuron types.^30^

### In L6, excitatory inputs are chiefly intralaminar

We next turned our attention to the spatial distribution of excitatory inputs onto L6 PCs, BCs, and MCs. L6 PCs and MCs received more excitatory inputs from L6 than inputs from the other layers, but L5 PC→L6 BC and L6 PC→L6 BC connectivity rates were indistinguishable (Figures 5A and 5B). L6 BCs also had overall higher excitatory connectivity rates from L5 than L6 PCs or L6 MCs did.

**Figure 5.**
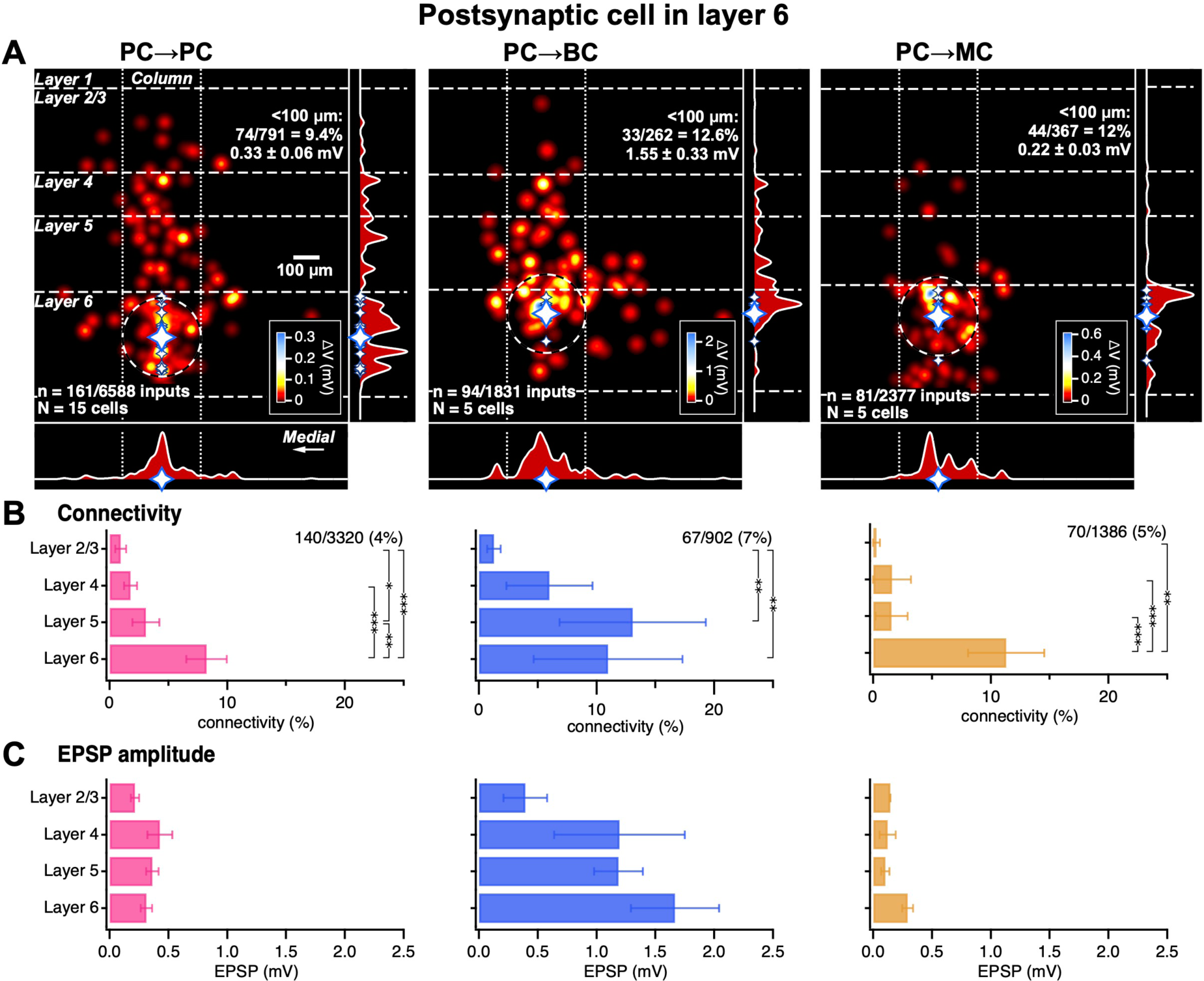
In L6, excitatory inputs to PCs, BCs, and MCs were chiefly from L6. (A) Synaptic input density maps for L6 PCs (left), BCs (middle), and MCs (right) were generated by averaging individual maps (Figure 1H). Insets right and bottom show the vertical and horizontal density projections, respectively. Connectivity within a 100-µm radius (dashed circle) enables comparison with paired patch-clamp recordings.^30^ Dashed horizontal lines: layer boundaries. Dotted lines: vertical column. (B) L6 PCs and MCs both received more excitatory inputs from L6 than from L2/3, L4, and L5. L6 BCs, however, received fewer excitatory inputs from L2/3 than from other layers, with an indistinguishable number of inputs from L5 and L6. BCs received more L5 inputs than did PCs or MCs (PC vs. BC, p < 0.01; BC vs. MC, p < 0.001). GLMM statistics revealed that connectivity depended on presynaptic layer and cell type (p < 0.001). (C) L6 BCs received stronger excitatory inputs than L6 PCs and L6 MCs, irrespective of the layer of origin of the input (PCs vs. BCs, p < 0.01, BCs vs. MCs, p < 0.01). LMM statistics revealed that EPSP amplitudes depended on cell type (p < 0.001) but not on presynaptic layer (p = 0.08, interaction effect p = 0.16). Mean ± SEM. \**P < 0.05*, \*\**P < 0.01*, \*\*\**P < 0.001*.

Within each cell type in L6, there was no difference in connection strength from different layers (Figure 5C). But across cell types in L6, excitatory inputs onto BCs were stronger than excitatory inputs onto PCs and MCs.

Taken together, these findings reveal relatively similar spatial distributions of excitatory inputs onto PCs, BCs, and MCs in L6. However, overall excitation of L6 BCs was appreciably stronger than that of L6 PCs or MCs. Since this was also true for BCs in L2/3 (Figure 3) and L5 (Figure 4), this suggests that stronger excitatory drive of BCs is a general principle. Also, we once again found that L6 was relatively isolated from the other layers, because — just like L6→L2/3 (Figure 3) and L6→L5 pathways (Figure 4) were weak — excitatory connectivity rates from outside L6 onto L6 were low.

### Excitation of inhibition originates farther away

We noticed from density maps that excitation of BCs and MCs (E→I) tended to draw on presynaptic PCs located farther away (Figures 3-5). To explore this idea, we quantified the distance-dependent decay of connectivity, *τ*, by fitting exponentials to radially binned connectivity rates (Figure S7; see Methods).

With this approach, we indeed found smaller values of *τ* for PCs than for BCs and MCs in L5 and L6, although this did not hold true for L2/3 (Figure S7), akin to L2/3 of mouse primary auditory cortex.^71^ In conclusion, E→I inputs originated more distally than excitation of PCs (E→E), at least for interneurons in subgranular L5 and L6. This could promote stability and difference-of-Gaussians connectivity.

### Optomapping and paired patch yield indistinguishable results

To validate optomapping, we compared it with L5 paired-recording data that we previously acquired.^46^ One caveat, however, is that our paired-recording study was carried out at a younger age range (paired recordings, P11-P20: 15.6 ± 0.13, n = 223 vs. optomapping, P17-P25: 21.3 ± 0.32, n = 41, p < 0.001, Wilcoxon-Mann-Whitney). Another is that paired-recording studies of the same brain region have been known to disagree,^30,31^ meaning this method is not a gold standard. Still, the comparison is useful, since paired recordings have long been the state of the art.

Our paired recordings were done with cells spaced as closely as possible, meaning <100 µm apart.^26^ To enable comparison, we therefore restricted the optomapping data set to within a 100-µm radius of the postsynaptic cell (Figure 4A, dashed circles).

For L5 PC→L5 PC paired recordings, we previously found^46^ a connectivity rate of 81/682 = ∼11.9% (see Methods), which was not different from the 15.5% of optomapping (Figure 4A, p = 0.084, chi-square). However, the L5 PC→L5 PC paired-recording EPSP amplitude (0.87 ± 0.09 mV, n = 162) was larger than that measured with optomapping (0.32 ± 0.03 mV, n = 64, p < 0.001, Wilcoxon-Mann-Whitney). However, the smaller optomapping amplitudes were in agreement with paired recordings from older mice,^30^ consistent with developmental downregulation of L5 PC→L5 PC release.^72^

For L5 PC→L5 BC synapses, connectivity rates obtained with paired recordings^46^ (100/299 = ∼33.4%) were indistinguishable from those acquired with optomapping (35.2%, Figure 4A, p = 0.67, chi-square). We were furthermore not able to tell apart L5 PC→L5 BC EPSP amplitudes acquired with the two methods (pairs, 2.10 ± 0.21, n = 100 vs. optomapping, 2.31 ± 0.41 mV, n = 82, p = 0.66, Wilcoxon-Mann-Whitney). Likewise, for L5 PC→L5 MC synapses, we found indistinguishable connectivity rates with paired recordings (4/47 = ∼8.5%) and optomapping (16.7%, Figure 4A, p = 0.17, chi-square). The L5 PC→L5 MC EPSP amplitudes were also indistinguishable with the two methods (paired recordings, 0.21 ± 0.1 mV, n = 4 vs. optomapping, 0.36 ± 0.08 mV, n = 27, p = 0.44, Wilcoxon-Mann-Whitney).

Overall, we found no consistent differences between paired recordings and optomapping. The weight differences for L5 PC→L5 PC synapses could be attributed to different animal ages.^72^ We concluded that results obtained with the two methods were indistinguishable.

### Excitatory pathway structure depends on target cell type

In V1, excitatory PC→PC pathways within and across layers have been well studied, which has led to the classical concept of the canonical cortical circuit.^8-11^ But because paired recordings are slow, the corresponding local circuits for PC→BC and PC→MC pathways across the cortical thickness have not been systematically explored.^30^ The high throughput of optomapping, however, enabled us to investigate this.

Based on ensemble optomaps (Figures 3 to 5), we constructed connectivity matrices to summarize excitatory pathways onto PCs, BCs, and MCs, which highlighted a prominent L4→L2/3 pathway for PCs (Figure 6A), as expected for V1.^8-11^ For BCs, connectivity was concentrated to L5 PC→L5 BC synapses. For MCs, however, the bulk of the connectivity was found at L2/3 PC→L2/3 MC connections.

**Figure 6.**
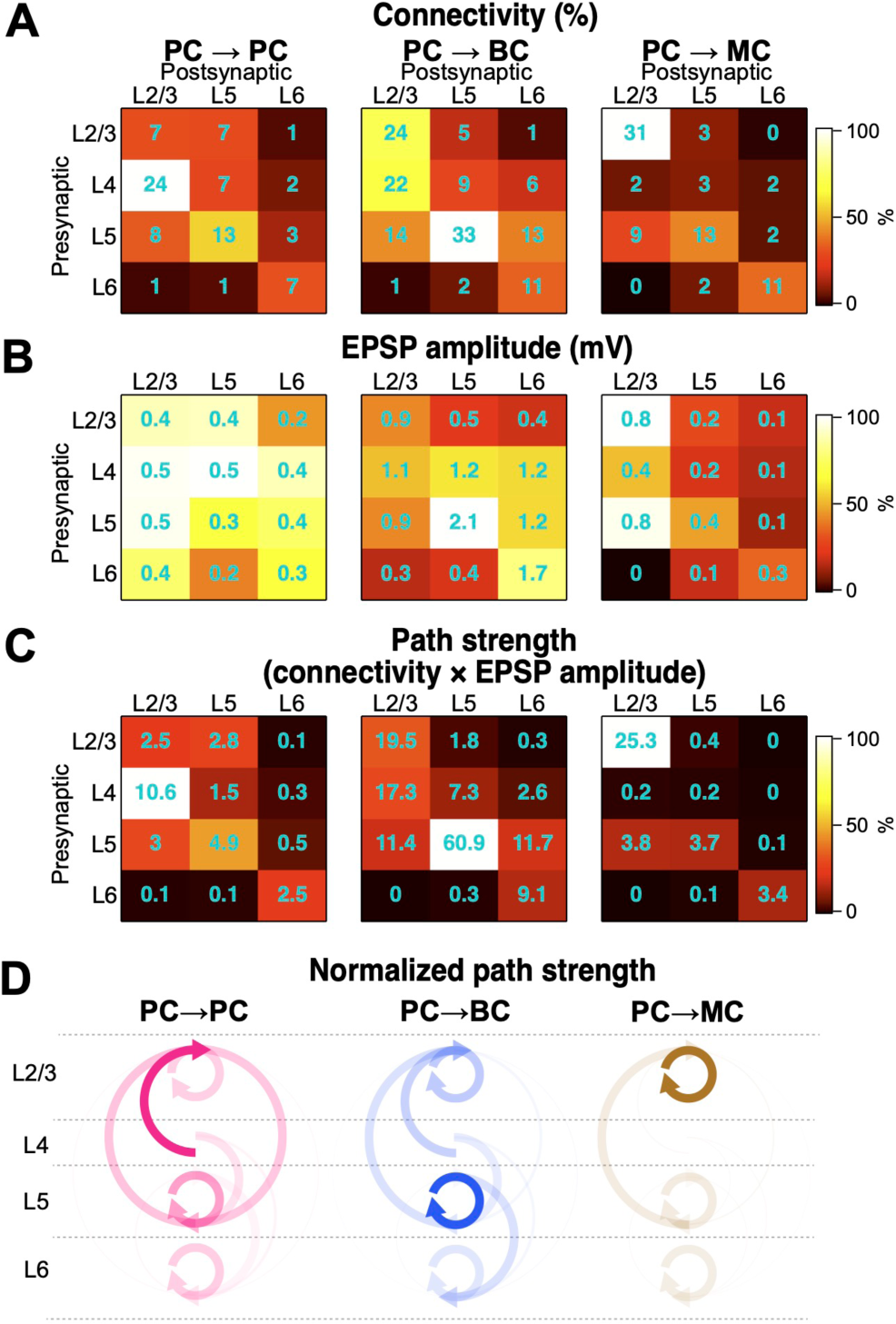
Excitatory circuit structure depends on target cell type. (A) Excitatory connectivity depended on pre- and postsynaptic layer in a target-cell-type specific manner. For PCs, L4→L2/3 connectivity was prominent. For BCs and MCs, however, L5→L5 and L2/3→L2/3 stood out, respectively. Color code denotes within-matrix normalized values, which highlight differences within a target cell type. Cyan numbers indicate absolute values, which enables comparison across target cell type. For statistical comparisons, see Figures 3 to 5. (B) Excitatory input strength onto PCs generally did not depend on input or output layer. For BCs, subgranular weights dominated, whereas for MCs, supragranular weights dominated. MC matrix shows EPSP1, so does not account for strong PC→MC facilitation (Figure S8). Color code, cyan numbering, and statistics as in (A). (C) Path strengths — i.e., the product of connectivity and EPSP amplitude^29^ — highlighted the strong L4 → L2/3 path for PCs, but for BCs, L5 → L5 was strongest, and for MCs, L2/3 → L2/3 was strongest. Overall excitation of PCs was weaker than excitation of either BCs or MCs. (D) Based on path strengths in (C), we constructed circuit schematics, which for PC→PC synapses reproduced the V1 canonical circuit.^10,11^ PC→BC paths, however, were strongest in L5, and PC→MC paths were strongest in L2/3. Arrow opacity indicates normalized pathway strength as shown in (C). Arrows on the left side indicate ascending paths and arrows on the right side denote descending paths. Circular arrows designate intralaminar pathways.

To explore if synaptic efficacy distributed similarly to synaptic connectivity, we created synaptic-strength matrices (Figure 6B). For PC→PC connections, synaptic efficacy distributed relatively evenly across the layers. For BCs, EPSP amplitudes were stronger in subgranular layers, especially for L5 PC→L5 BC synapses. In contrast, for MCs, EPSP amplitudes dominated in supragranular layers. Synaptic weight patterns for BCs and MCs (Figure 6B) thus echoed the differential connectivity rates (Figure 6A).

To investigate the combined effect of connectivity and synaptic efficacy, we created path strength matrices (Figure 6C), where path strength is defined as the product of connectivity and EPSP amplitude.^29^ For PCs, this matrix highlighted the strong L4→L2/3 path. For BCs, however, L5→L5 intralaminar drive was prominent. For MCs, on the other hand, L2/3→L2/3 intralaminar drive was more salient. Overall, path strengths onto PCs were weaker.

In summary, we found that excitatory microcircuit structures were target cell specific (Figure 6D). For PCs, we reproduced the V1 canonical circuit,^8-11^ but we found surprising differences in excitatory input structure for BCs and MCs. Finally, E→E pathways were on average weaker than E→I pathways.

### Short-term plasticity depends on layer as well as target cell type

It is known that short-term plasticity is specific to target cell,^15^ e.g., PC→MC synapses short-term facilitate but PC→BC connections short-term depress.^16,17,46,69^ Similarly, neocortical PC→PC connections generally short-term depress.^16,17,46,69^ However, given the distinct computational roles of the different cortical layers^1,2,8^, we wondered if these short-term plasticity patterns apply to all cortical layers, as indicated by paired recordings.^30^

Prior studies indicate that direct 1-photon optogenetic stimulation of synaptic terminals can artificially skew synaptic dynamics toward short-term depression by directly depolarizing and increasing calcium influx into presynaptic terminals.^73-75^ We therefore wanted to first verify that optomapping correctly identified short-term plasticity. We simultaneously optomapped excitatory inputs onto a PC, a BC, and an MC (Figure 7A), which qualitatively revealed target-cell-specific synaptic dynamics even for the same presynaptic PCs (Figure 7B).^16,17,69^ Quantification of short-term plasticity with paired-pulse ratio (PPR) revealed that synaptic responses in the PC short-term depressed, those in the BC less so, while those in the MC short-term facilitated (Figure 7C). This agreed with prior reports that target cell type determines short-term plasticity,^16,17,46,69^ demonstrating that optomapping accurately identifies synapse-type-specific synaptic dynamics. We furthermore detected no bias toward short-term depression,^73-75^ as expected from the soma-targeted ChroME expression and photo-stimulation. Finally, there was a good correspondence between optomapping and paired-recording PPR values for PCs, BCs, and MCs in L5.^46,76^

**Figure 7.**
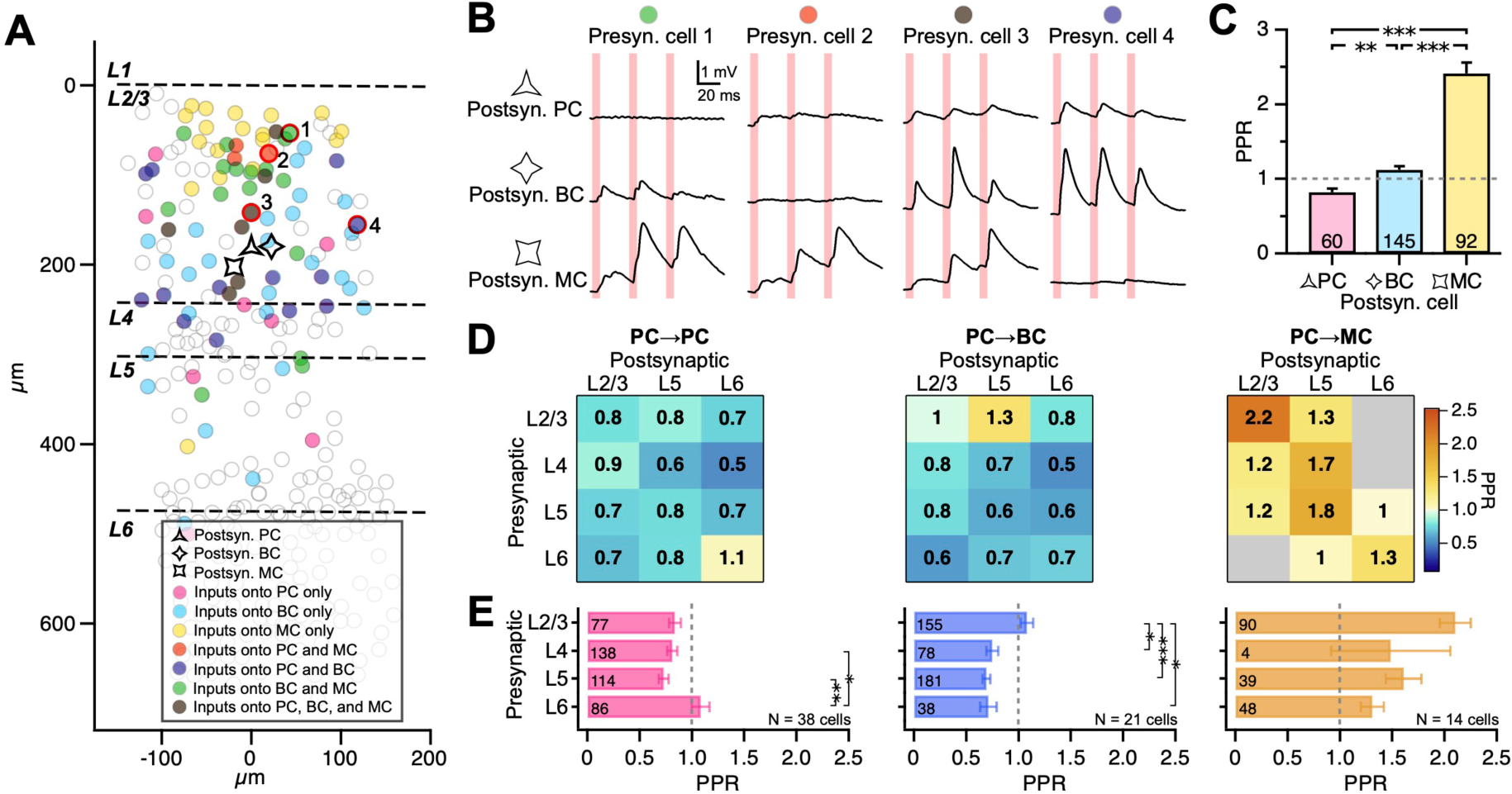
Synapse dynamics varied with cortical layer as well as target cell type. (A) Sample experiment where a PC, a BC, and an MC were simultaneously optomapped (same as in Figure S2). We found shared inputs onto all three postsynaptic cells. For clarity, inputs from other z-axis planes are not shown (Figure S2). (B) For the optomap in (A), presynaptic cells 1, 2, 3, and 4, denoted by red circles in (A), elicited EPSPs in more than one postsynaptic cell. However, the same presynaptic PCs elicited different short-term plasticity depending on target cell type, e.g., with short-term depression onto the PC and short-term facilitation onto the MC. EPSP traces are averages of 20 repeats. Pink bars denote spiral scan duration. (C) Across all connections optomapped in (A), PC→PC connections short-term depressed, PC→BC less so, and PC→MC synapses facilitated strongly, revealing that synaptic dynamics depended on target cell type, as previously shown.^16,17,46,69^ Dotted grey line demarcates short-term facilitation from short-term depression. (D) PPR measurements across different pre and postsynaptic layers for PCs, BCs, and MCs reinforced the principle that target cell type determines short-term plasticity patterns.^15^ However, apparent outliers included L6 PC→PC, L2/3 PC→L5 BC, L5 PC→L6 MC, and L6 PC→L5 MC connections. Overall, an additional dependence of synaptic dynamics on pre and postsynaptic layer emerged. LMM statistics revealed that PPR depended on presynaptic layer and postsynaptic cell type (p < 0.05), but not postsynaptic layer (p = 0.12), see (E). For inputs from L2/3 and L5, MCs had greater PPR than PCs and BCs (PCs vs. MCs, p < 0.001, PCs vs. BCs, p < 0.001). For inputs from L6, postsynaptic BCs had lower PPR than PCs and MCs (PCs vs. BCs, p < 0.01, BCs vs. MCs, p < 0.001). Numbers indicate mean PPR. Grey: category with fewer than three connections (see Methods). (E) As revealed in (D) and by LMM statistics, synaptic dynamics of inputs onto PCs and BCs showed an additional dependence on presynaptic layer location. L6 PC→PC inputs showed greater PPR than L4 and L5 PC→PC inputs. L2/3 PC→BC inputs showed greater PPR than L4, L5, and L6 PC→BC inputs. Since PPR did not depend on postsynaptic layer, we pooled across postsynaptic layers L2/3, L5, and L6 for PCs, BCs, and MCs. Numbers in each bar indicate number of inputs. Dotted grey line demarcates short-term facilitation from short-term depression. Mean ± SEM. *\*P < 0.05,* \*\**P < 0.01*, \*\*\**P < 0.001*.

To systematically explore target-cell-specific short-term plasticity across the cortical thickness, we broke the PPR measurements down with respect to presynaptic layer, postsynaptic layer, and postsynaptic cell type (Figure 7D). This revealed predominant short-term depression for PC→PC and PC→BC connections. However, noteworthy exceptions to the rule included L6 PC→PC and L2/3 PC→L5 BC pathways, which facilitated weakly. PC→MC synapses, on the other hand, consistently facilitated, although with variability across layers (Figure 7D).

We found that PPR did not depend on the layer location of the postsynaptic cell, so for statistical comparisons, we pooled data for inputs from the same presynaptic cell layer onto each cell type (Figure 7D). This revealed stronger facilitation from L6 inputs for PCs but stronger facilitation from L2/3 inputs for BCs (Figure 7E). For MCs, however, short-term dynamics were indistinguishable across layers (Figure 7E), suggesting high heterogeneity (Figure 7D).

We were with optomapping thus able to produce an excitatory short-term plasticitome for PCs, BCs, and MCs in developing V1.^13,35^ This revealed that synaptic dynamics depended on cortical layer and not just on target cell type,^15^ suggesting complex interactions between pre- and postsynaptic partners.

### Excitation latency varies with target cell type

Excitatory inputs onto MCs show strong short-term facilitation while those onto PCs and BCs short-term depress (Figure 7).^23,30,46^ In addition, MCs have a long membrane time constant compared to BCs (Figure S4C),^46^ leading to appreciable temporal summation in MCs for the 30- Hz EPSP trains we evoked (Figure 1D). Therefore, relying on the first response, EPSP_1,_ in a train of three underestimates excitation of MCs.

We measured the peak depolarization, ΔV_peak,_ due to the temporal summation of the three EPSPs. and we found that the excitation of MCs was as expected now markedly stronger (Figure S8A and S8B). Moreover, using ΔV_peak t_o construct circuit maps brought out the weakness of PC→PC synapses relative to PC→BC and PC→MC connections while emphasizing the differential circuit structures we already found (Figures 6C and S8C).

It is known that, because of their different excitatory input short-term dynamics, BCs fire early during a high-frequency burst and thereby mediate fast-onset inhibition^21,22^ whereas MCs fire with a delay, to provide late-onset inhibition.^21,23^ Consequently, we argued that peak depolarization should serve as a meaningful proxy for the latency at which postsynaptic neurons tend to fire. As expected, BC latencies were short^21,22^ and MC latencies were long^21,23^ (Figure S8D). Surprisingly, however, PC latencies distributed heterogeneously (Figure S8D).

Overall, these findings highlighted the long latency of peak excitation of MCs. This observation reinforced the principle that E→E pathways are generally weaker than E→I projections. Finally, the heterogeneity of latencies in PCs suggests that they are sensitive to changes in excitatory input short-term dynamics.^77-79^

### High-order connectivity patterns in L6

Classical connectivity studies have revealed high-order connectivity patterns in L2/3 and L5.^26,32,33,64,65^ We therefore examined L6 for high-order structure. To do so, we combined data from optomapping and multiple patch. We relied on Monte-Carlo bootstrap analysis (see Methods) of optomapped L6 PC pairs to essentially carry out in silico the equivalent to paired-patch connectivity sampling in local circuits (Figure 8A-8C).^26,32,33^ This way, we estimated that reciprocally connected L6 PC↔PC pairs had a four-fold overrepresentation of shared excitatory inputs, whereas the overlap of shared inputs for unconnected L6 PCs (N = 3 pairs pooled) was indistinguishable from uniformly random (Figure 8D). Whereas several prior studies show high-order connectivity pooled data across postsynaptic cells,^26,32,33^ we show that such patterns hold true even for individual neuronal pairs.

**Figure 8.**
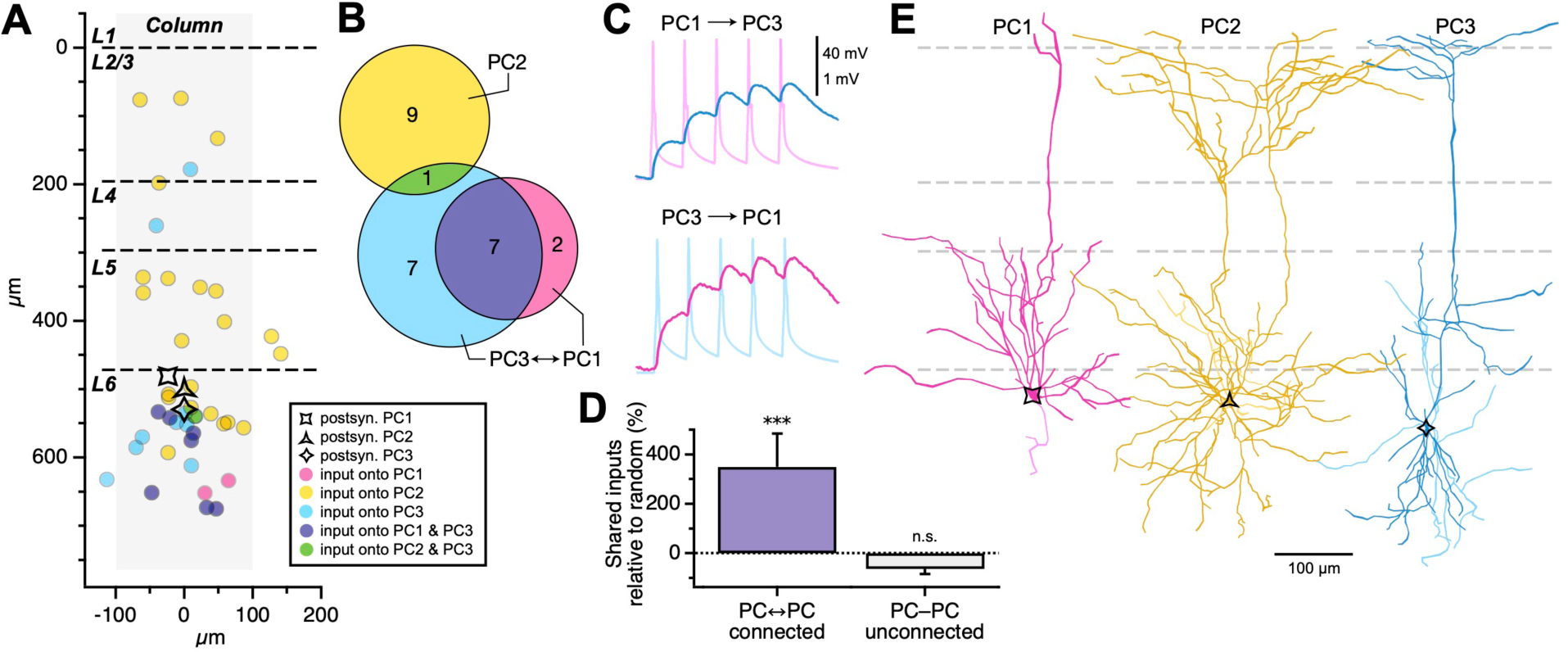
Four-fold overrepresentation of shared inputs onto connected L6 PCs. (A) Sample experiment where three L6 PCs were simultaneously optomapped and inputs onto each of the PCs were identified. PC2 received inputs throughout the height of the column, while inputs onto PC1 and PC3 were generally restricted to L6. For clarity, only inputs that elicited EPSPs are shown. (B) Of the inputs identified in (A), we identified several that were shared by two of the patched PCs. We did not observe any inputs shared by all three PCs. (C) PCs 1 and 3 were bidirectionally connected. To determine connectivity across patched PCs, we elicited 5 spikes at 50 Hz in each PC by sequential current injections while recording from the other PCs. EPSP traces are averages of 20 repeats. (D) Reciprocally connected PC pair shared more L6 inputs than expected from uniformly random, as revealed by Monte-Carlo bootstrap (see Methods). The number of L6 inputs shared by PCs that were not connected (N = 3 pairs pooled) were indistinguishable from uniformly random. (E) Morphological reconstruction verified that all three cells were pyramidal. Although PC2 appeared more branched, this distinction did not hold true for the other unconnected L6 PCs (not shown). Dark lines: dendrite; lighter lines: axon. \*\*\**P < 0.001*

We wished to validate our optomapping approach by reproducing a previously published result. It was previously demonstrated that reciprocally connected L2/3 PC↔BC pairs share more excitatory inputs than expected from uniformly random, but that this is not true for PC-MC pairs, regardless of within-pair connectivity.^65^ Applying the same bootstrap approach to the multiple-patch data in Figure 7, we indeed found a ∼30% overrepresentation of shared inputs onto reciprocally connected L2/3 PC↔BC pairs (excess relative to uniformly random µ ± SDev = 34.3% ± 18%, p < 0.01, Monte-Carlo bootstrap, data not shown). However, the equivalent overrepresentation was not found for the unconnected L2/3 PC-MC pair (-22.4% ± 17%, p = 0.87, Monte-Carlo bootstrap). This outcome agrees with the prior literature,^65^ lending support to our approach.

In sum, we found that interconnected L6 PCs shared an excess of excitatory inputs, whereas unconnected L6 PCs did not. The existence of high-order connectivity patterns is thus a general principle that extends from L2/3 and L5 to L6. This principle furthermore also holds true for individual neuronal pairs.

## DISCUSSION

Our study showcases optomapping, a high-throughput 2P optogenetic synapse-charting approach that we validate as accurate and reliable. Many classic studies explored cortical pathways in bulk, whereas optomapping permitted us to rapidly interrogate a synapse at a time. We were therefore able to reveal several hitherto unappreciated principles that govern V1 excitatory fine structure, such as synaptic-weight distribution log-normality in single cells, target-cell-specific circuit structure, and layer-dependent synaptic dynamics.

### The ubiquitous log-normal distribution

Many physiological and anatomical features in the brain are described by long-tailed power-law or log-normal distributions.^34,80^ For instance, spine size distributes log-normally,^81^ and since spine size scales with synaptic strength,^80,81^ this implies that weight distributions should be similarly heavy-tailed. In agreement, paired recordings have revealed that unitary synaptic responses distribute log-normally.^26,29,34,52^ But because paired-recording studies pool data across cells,^26,29,52^ log-normal weight distributions could arise artificially due to cross-cell sampling. For example, most cells may be weakly driven, but hub neurons may receive numerous, strong inputs.^80^ However, we found log-normal distribution in individual PCs (Figure 2), meaning it could not have arisen from cross-cell sampling.

Interneurons, on the other hand, are largely devoid of spines,^18-20^ so spine size says little about synaptic weights onto BCs and MCs. The weight distribution of interneurons has thus been unclear.^34^ Log-normal weight distribution means few synapses elicit strong responses, whereas most are vanishingly weak. This long-tailed weight distribution is important because may underlies key functional capabilities such as feature encoding.^28^ Consequently, PCs that code for similar information interconnect more often and more strongly,^27^ as expected from Hebbian plasticity binding these PCs together. In the same vein, it has been proposed that log-normal synaptic strengths emerge from Hebbian-like plasticity,^53^ yet many interneuron types do not exhibit Hebbian plasticity.^13^ In particular, PC→BC synapses undergo anti-Hebbian plasticity.^54,55^ We therefore speculated that excitatory weight distributions of BCs might not be log-normal.

However, we found log-normality in both BCs and MCs, even in individual interneurons (Figure 2). Our results thus further cemented log-normality as a ubiquitous feature of synapses formed by PC axons, although the precise mean and standard deviations varied with target cell type. It is unclear if axons of inhibitory cells also generally produce synapses with log-normally distributed weights, but pooling across cells suggests so.^68^

Theory studies have suggested that heavy-tailed distributions can arise from multiplicative processes such as homeostatic, intrinsic, or structural forms of plasticity.^53,82,83^ The causative agent of log-normality thus remains unclear, although our BC findings do not favor Hebbian plasticity. Interestingly, experiments show that spine size log-normality emerges even under synaptic blockade,^81^ suggesting activity-dependent plasticity is not required.

Finally, we reproduced in single cells the known sparsity of cortical connectivity (Figures 3 to 5),^26,29,52^ with most weights being zero.^34,84^ It has been shown that the existence of a large fraction of zero weights — i.e., potential synapses as a blank slate^5^ — is key to optimal information storage in cortical circuits.^84^

### The structure of excitatory circuits

Neocortical circuits generalize across areas, repeating the same basic laminar organization of neocortical excitatory neurons.^10^ In a simplistic view, this canonical circuit consists of an ascending path from L4 to L2/3, the computational layer, which then projects to L5, the output layer.^10,11^ However, classic studies typically explored these pathways in bulk, whereas optomapping permitted the study of canonical circuits a synapse at a time, enabling interrogation of sublaminar structure.

PC→PC optomapping largely reproduced the classical canonical circuit, although we found intriguing differences compared to prior literature. It has been reported that L2/3→L5 is stronger than the L2/3→L2/3 projection,^67^ but we found that these paths were indistinguishable. Like others,^68^ we additionally found a prominent L5→L2/3 projection, yet this pathway is absent from influential working models of neocortical circuits,^85,86^ as well as from several paired recording studies.^29,31,72^ Another striking observation with optomapping was the relative isolation of L6 from the other cortical layers, as shown before,^30^ although the documented L5→L6 projection^10,11^ did show up weakly. L6 may thus be relatively computationally independent, although in the intact brain, neuromodulation may turn on pathways to and from this layer.

Key cortical interneuron types such as BCs and MCs establish functionally important recurring connectivity motifs.^21-25^ We therefore considered BCs and MCs as targets in addition to PCs, which revealed striking differences (Figure 6). PC→BC excitation was overwhelmingly concentrated to L5. PC→MC excitation, on the other hand, dominated in L2/3, reminiscent of barrel cortex.^70^ In both cases, there was an intriguing preponderance for excitation to originate in the upper half of the layer (Figures 3 and 4), suggesting sublaminar structuring.

So, although PC→PC optomapping chiefly reproduced the known canonical circuit with some fine-tuning, the layer-specificity of PC→BC and PC→MC drive was surprising. For example, MCs have been thoroughly studied in L5,^21,23,24,46,51,76^ yet are much more strongly driven in L2/3, hinting at a key functional relevance in the computational layer. Similarly, the potent PC→BC drive in L5 suggests a prominent BC role in the output layer.

### E→I: denser, stronger, farther

Throughout this study, we noticed that E→I synapses were both denser and stronger than E→E synapses (Figures 3 to 6). Both connectivity and EPSP amplitudes were thus stronger for PC→BC and PC→MCs than for PC→PC synapses, which was true across all layers. This differential outcome for E→E versus E→I drive was particularly striking once we accounted for the strong facilitation of PC→MC connections (Figure S8).^15,21,23^

In addition, excitatory inputs onto BCs and MCs generally had a greater spatial reach than did synapses onto PCs (Figure S7). This difference, however, was not true for L2/3, similar to L2/3 of mouse primary auditory cortex.^71^ It is also possible that the strong ascending L4→L2/3 pathway (Figure 3) distorted exponential fits specifically for L2/3 PCs.

Overall, an emerging principle is nevertheless that E→I projections tend to be denser, stronger, and farther-reaching than E→E projections. This differential arrangement of E→E and E→I synapses may prevent runaway excitation in local circuits since local hyperactive PCs are efficiently and promptly shut down due to their strong drive of neighboring inhibitory BCs and MCs.

The combination of the E→I and E→E radial distributions furthermore provides a substrate for Mexican-hat-shaped difference-of-Gaussian connectivity structures.^87^ It has long been argued that such spatial connectivity patterns mediate lateral suppression in local circuits and underlie edge detection in vision.^88^ Likewise, L2/3 I→E projections are relatively long-range,^68^ lending support to this idea.

### A V1 excitatory short-term plasticitome

It is a well-known principle that short-term plasticity is synapse-type specific.^15^ For instance, PC→BC connections short-term depress but PC→MC synapses facilitate, which is one reason why BCs and MCs elicit early and late-onset inhibition, respectively.^16,17,46,69^ Since the layers are known to have distinct computational roles,^1,2,8^ we wondered if this principle held true across the V1 thickness. By optomapping the short-term dynamics of PC→PC, PC→BC, and PC→MC synapses across the cortical lamina, we therefore created an excitatory short-term plasticitome (Figure 7).^13^

As expected, PC→MC synapses facilitated and PC→PC as well as PC→BC synapses generally short-term depressed. This verified the known target-cell specificity of synaptic dynamics,^15^ which in turn validated optomapping as a reliable high-throughput method for measuring short-term dynamics.

However, we additionally found layer-specific differences, e.g., stronger facilitation in L6 inputs for PCs and stronger facilitation in L2/3 inputs for BCs. To our knowledge, these cell-specific dissimilarities across the lamina have not been previously reported.^30,31^ Our finding suggests that, over the course of cortex-wide sustained activity, E→E and E→I drive will vary differentially in supra and subgranular layers. In other words, late inhibitory spiking may dominate in the top layers, but late excitatory spiking may dominate in the bottom layers, suggesting an excitatory wave of information from pia toward white matter during bursting activity.

A most parsimonious interpretation is that E/I balance also varies accordingly across the cortical thickness. Verifying this interpretation, however, would require also optomapping I→E projections, which we did not investigate here, but which are possible to optomap.^68^

Factors other than synaptic short-term dynamics also determine BC and MC spiking latency, such as membrane time constant and time course of synaptic conductances.^76^ To account for all such factors collectively, we also explored temporal summation in the form of peak depolarization latency (Figure S8). Unsurprisingly, depolarization peaked early for BCs and late for MCs, as expected from their known roles in early and late-onset inhibition.^16,17,46,69^

Unexpectedly, peak depolarization latencies in PCs distributed relatively heterogeneously over short EPSP bursts. PC→PC connections are well known for undergoing presynaptic forms of long-term plasticity that alter short-term synaptic dynamics.^77-79^ For presynaptically expressed long-term plasticity, long-term potentiation redistributes synaptic efficacy toward the beginning of high-frequency EPSP trains,^78,79^ whereas long-term depression shifts synaptic weight toward the end.^77^ Given the relatively heterogenous temporal distribution of peak depolarization latencies in PCs, presynaptically expressed long-term plasticity at PC→PC connections is thus well positioned to alter not just the likelihood of postsynaptic PC spiking, but also its latency relative to that of BCs and MCs. This is a previously unappreciated perspective on E→E plasticity that contextualizes it in the plasticitome of the local circuit by contrasting with E→I and I→E projections.^13^ Testing and developing this view would require computer models of cortex-wide long-term plasticity that include the pre-versus postsynaptic locus of expression of plasticity,^89,90^ which is outside the scope of the present study.

### High-order patterns of connectivity

Several influential connectivity studies have revealed the existence of high-order connectivity patterns.^26,32,33,64,65^ For instance, compared to unconnected L5 PC pairs, reciprocally connected L5 PCs more likely receive input from the same L2/3 PC.^32^ High-order microcircuit structure is important as it shapes information processing in the brain, e.g., by binding different features of information or by creating separate streams of information.^26,32,33,64,65^ However, many of these studies were carried out across experiment days and animals with paired recordings.^26,32,33^ They also focused on L2/3 and L5.^26,32,33,64,65^

To cover new ground, we therefore focused on L6, where we explored if this connectivity principle carried over to individual PC pairs. We found a striking four-fold overrepresentation of shared inputs onto connected L6 PCs that was not present for unconnected L6 PCs (Figure 8). To validate our finding, we also reproduced prior findings on high-order connectivity in L2/3.^65^

Patchy high-order connectivity patterning has long been reported in L2/3 and L5,^26,49,63^ which has in some cases been attributed to the existence of different PC types,^12,33,91^ and in other cases not.^32,64^ We were not able to independently classify unconnected from connected L6 PCs (Figures 8E, S4). Although we could not find evidence that our high-order connectivity patterns necessarily aligned with L6 PC type, we also could not exclude this possibility. Overall, our findings agreed with prior literature on V1 L6 PCs,^92,93^ which supports the view that local L6 PC connectivity relates to L6 PC projection patterning.

The existence of high-order connectivity patterns thus extends to L6, even for individual pairs. Furthermore, optomapping is a reliable and efficient method for assessing high-order connectivity patterns.

### Outlook

The state-of-the-art approach for synapse-type-specific experimentation has long been the paired-recording technique.^26,27,29-33^ This methodology, however, is difficult to learn and slow to use, leading to the throughput problem.^13,35^ Consequently, relatively complete mappings of entire microcircuits have been rare.^29-31^ Since circuit structure fundamentally determines circuit function,^1-3^ the throughput problem has therefore been a major impediment to progress in neuroscience research.

To solve the throughput problem, we created optomapping by seeking inspiration from recent advances with 2P optogenetics.^37,68,94^ We validated optomapping by comparing with paired recordings. We could with optomapping rapidly and reliably test hundreds of candidate inputs hundreds of microns away from a patched cell, across the cortical layers and covering the entire cortical thickness, to reveal hitherto unappreciated microcircuit differences for PCs, BCs, and MCs. We estimate that optomapping is around two orders of magnitude faster than multiple patch. For even better yield, optomapping can also be combined with other approaches, such as multiple patch^26,27,29-31^ and patch robots.^95,96^

Optomapping was not without caveats. We identified direct optogenetic activation of opsin-expressing postsynaptic cells as a central problem as well as approaches to mitigate it. Ideally, however, opsins in postsynaptic cells should be specifically blocked by drug dialysis via the recording pipette.^46,51^ Development of such pharmacology would be a key step forward for optomapping reliability.

Because of the throughput problem, the typical medium-sized lab has not been able to explore how local circuits differ in disease states, in genetic models, across brain areas, or across species. By solving this problem, optomapping thus changes what kinds of questions neuroscientists can ask. Here, we showcased optomapping by applying it to developing V1, which provided fresh perspective on the principles that govern its excitatory fine structure.

## MATERIALS AND METHODS

See supplemental information for details.

## SUPPLEMENTAL INFORMATION

Document S1. STAR Methods, Figures S1-S8, and Supplemental References.

## ACKNOWLEDGMENTS

We thank Alanna Watt, Charles Bourque, Keith Murai, Aparna Suvrathan, Arjun Krishnaswamy, Ed Ruthazer, Jonathan Britt, Jon Sakata, Amanda McFarlan, Haley Renault, Shawniya Alageswaran, Connor O’Donnell, Christopher Salmon, W. Todd Farmer, Riccardo Beltramo, Or Shemesh, Yoaz Printz, Ofer Yizhar, Hillel Adesnik, Adam Packer, Karl Deisseroth, and Sjöström lab members for help and useful discussions.

C.Y.C.C was in receipt of a McGill University Max E. Binz Fellowship, Healthy Brains for Healthy Lives (HBHL) PhD Fellowship, NSERC CGS D 534171-2019, FRNTQ B2X 275075, and the Ann and Richard Sievers Neuroscience Award. H.H.-W.W. was supported by CIHR Fellowship 295104, CFREF, HBHL Postdoctoral Fellowship, FRSQ Postdoctoral Fellowship 259572, and QBIN Scholarship 35450. C.G. won NSERC USRA, FRQNT BPC, and RI-MUHC studentships. K.E.B. was funded by an IBRO-Brain Studentship. T.L. was awarded an NSERC USRA. P.J.S. acknowledges funding from CFI LOF 28331, CIHR PG 156223, FRSQ CB 254033, and NSERC DG/DAS 2017-04730 as well as 2017-507818. The Montreal General Hospital Foundation kindly funded the Chameleon ULTRA II laser.

## AUTHOR CONTRIBUTIONS

Conceptualization, C.Y.C.C. and P.J.S.; methodology, C.Y.C.C. and P.J.S.; investigation – optomapping, C.Y.C.C.; investigation – quadruple patch clamp, C.Y.C.C. and H.H.-W.W.; investigation – neuronal reconstructions, C.Y.C.C., K.E.B., C.G., C.H., J.J., T.K., V.Y.L., T.A.L., and V.C.W.; custom software, P.J.S.; formal analysis, C.Y.C.C. and P.J.S.; writing – original draft, C.Y.C.C. and P.J.S.; writing – review & editing, C.Y.C.C., H.H.-W.W., and P.J.S.; funding acquisition, C.Y.C.C., H.H.-W.W, and P.J.S; supervision, P.J.S.

## INCLUSION AND DIVERSITY

We worked to ensure sex balance in the selection of non-human subjects. One or more of the authors of this paper self-identifies as an underrepresented ethnic minority in their field of research or within their geographical location. One or more of the authors of this paper self-identifies as a gender minority in their field of research. One or more of the authors of this paper received support from a program designed to increase minority representation in their field of research. While citing references scientifically relevant for this work, we actively worked to promote gender balance in our reference list.

## RESOURCE AVAILABILITY

Lead contact

Further information and requests for resources and reagents should be directed to and will be fulfilled by the lead contact, P. Jesper Sjöström.

Materials availability

This study did not generate new unique reagents.

Data and code availability

- All data reported in this paper will be shared by the lead contact upon reasonable request.
- All custom code used for data acquisition and data analysis are available from the lead contact on reasonable request, as well as in GitHub links provided in the Key Resource Table section.
- Any additional information required to reanalyze the data reported in this paper is available from the lead contact upon request.

## EXPERIMENTAL MODEL AND SUBJECT DETAILS

### Animals

Procedures were carried out in accordance with the *Canadian Council on Animal Care* and overseen by the Montreal General Hospital *Facility Animal Care Committee*, with appropriate licenses. Animals were kept on a 12 h:12 h light/dark cycle. Females and males were used for experiments. Homozygous Emx1^Cre/Cre^ mice^38^ were obtained from Jackson Laboratory, Bar Harbor, ME (strain number 005628).

## METHOD DETAILS

### Viral Infection

Adeno-associated viruses (AAVs) AAV9-CAG-DIO-ChroME-ST-P2A-H2B-mRuby3 (108912, Addgene, Watertown, MA) and AAV9-mDlx-GFP-Fishell-1 (83900, Addgene) were diluted to a titer of ∼2.7e12 GC/mL in sterile phosphate buffer solution (PBS) (Thermo Fisher Scientific, Waltham, MA) and aliquoted. Aliquoted AAVs were kept at -80°C until use. Intracerebral AAV injections were carried out in neonatal (P0-P2) Emx1^Cre/Cre^ mice^38^ according to previously published descriptions,^40^ although for higher expression levels, we injected directly into V1 rather than into the lateral ventricles. The two AAVs were mixed at a 2:1 ratio of ChroME to mDlx. Mice were cryoanesthetized and placed in a stereotaxic frame. The head was leveled on the anterior-posterior and medial-lateral axes, using bregma and skull landmarks, and fixed in place with the rubber-covered blunt end of ear bars. Using a 33-gauge needle attached to a 10 µL gas-tight syringe (Hamilton Instruments, Reno, NV), one V1 site was injected, 0.00 mm anterior-posterior, 1.05-1.15 mm medial-lateral, with respect to lambda suture coordinates. Three infusions of 0.2-0.3 µL were performed at depths of 0.20 mm, 0.15 mm, and 0.10 mm below the pial surface, at a rate of 0.25 µL/minute. To reduce backflow of AAV mixture upon needle retraction, the syringe was slowly removed over 2 minutes. Anesthetized mice were allowed to recover movement on a heating pad and subsequently returned to their home cage.

### Acute Brain Slice Electrophysiology

Injected mice aged P18-P25 were anaesthetized with isoflurane and decapitated once the limb withdrawal reflex was lost. Mouse brains were dissected in ice-cold (∼4°C) artificial cerebrospinal fluid (ACSF, in mM: NaCl, 125; KCl, 2.5; MgCl_2,_ 1; NaH_2P_O_4,_ 1.25; CaCl_2,_ 2; NaHCO_3,_ 26; Dextrose, 25; bubbled with 95% O_2/_5% CO_2)_. Osmolality of the ACSF was adjusted to 338 mOsm with glucose. Three-hundred-micron-thick near-coronal slices were cut from V1 with a Campden Instruments 5000mz-2 vibratome (Campden Instruments, UK), according to standard procedures.^97^ Slices were transferred to an incubation chamber, kept at 36°C for ten minutes, and then allowed to cool to room temperature (∼23°C). Experiments were carried out with ACSF heated to 33°C with a resistive inline heater (Scientifica Ltd, UK), with temperature recorded and verified off-line. Recordings were truncated or not used if temperature fell outside the 32-34°C range.

Patch pipettes of 4-7 MΩ resistance were pulled from medium-wall capillaries using a P-1000 electrode puller (Sutter Instruments, Novato, CA). Pipettes were filled with internal solution consisting of (in mM): KCl, 5; K-Gluconate, 115; K-HEPES, 10; MgATP, 4; NaGTP, 0.3; Na-Phosphocreatine, 10; and 0.1% w/v Biocytin, adjusted with KOH to pH 7.2-7.4 and with sucrose to 310mOsm. Internal solution was supplemented with Alexa Fluor 594, 10-40 µm, or Alexa Fluor 488, 80-120 µM (Life Technologies, Waltham, MA).

Whole-cell recordings were obtained using BVC-700A amplifiers (Dagan Corporation, Minneapolis, MN). Current clamp recordings were filtered at 5-6 kHz and acquired at 40 kHz using PCI-6229 boards (NI, Austin, TX) with custom software^77^ (available at https://github.com/pj-sjostrom/MultiPatch.git) running in Igor Pro 8 or 9 (WaveMetrics Inc., Lake Oswego, OR) on a custom-built computer (SL-DK-WS-PD-C236SAE-IF upgraded to quad-core Intel Core i7-6700, SuperLogics, Natick, MA). Series resistance, perfusion temperature, input resistance, and resting membrane potential, were monitored online and assessed offline (see below). Series resistance was not compensated. Liquid junction potential (10 mV) was not accounted for.

Neurons were patched with a LUMPlanFL N 40×/0.80 objective (Olympus, Olympus, Melville, NY) using infrared video Dodt contrast on a custom-modified Scientifica SliceScope.^46^ V1 was targeted based on the location of the dorsal hippocampal commissure white matter tract and proximity to the secondary visual cortex, which is distinct because it lacks L4. PCs were targeted based on their triangular shape and prominent apical dendrite. INs were targeted based on GFP expression visualized with 2P microscopy at 920 nm (see 2P Microscopy section below). Cell identity was verified *post hoc* using electrophysiological and morphological properties (see Cell Classification section below and Figures S4 and S5).

### Pharmacology

To assess 2P optogenetic activation of ChroME-expressing neurons (Figure S3), we blocked synaptic transmission using 200 µM D/L-AP5 and 5 µM NBQX (Hello Bio, Bristol, United Kingdom).

### 2P Microscopy

2P microscopy as well as 2P optomapping (see below) was performed with workstations custom-built from Scientifica SliceScope microscopes, as previously described.^46^ 2P excitation was achieved using a Chameleon ULTRA II (Coherent, Santa Clara, CA) titanium-sapphire laser tuned to 1040 nm, manually attenuated with a polarizing beam splitter in combination with a half-lambda plate (GL10-B and AHWP05M-980, Thorlabs Inc, Newton, NJ). Laser power was monitored by letting a glass slide reflect a fraction of the beam into a power meter (PM100A with S121C, Thorlabs). The laser beam was doubly gated with a shutter (SH05/SC10, Thorlabs) and a galvanometric mirror (GVS011/M, Thorlabs), the former for laser safety, the latter for speed and reproducibility. A pair of 6215H 3-mm galvanometric mirrors (Cambridge Technology, Bedford, MA) were used as beam scanners. Detectors were based on R3896 bialkali photomultipliers (Hamamatsu, Bridgewater, NJ) in a substage configuration. Fluorescence was collected with an FF665 dichroic and an FF01-680/SP-25 emitter (Semrock Inc., Rochester, NY). Red and green fluorescence were separated with an FF560-Di01 dichroic beam mirror (Semrock), a ET630/75 m (Chroma Technology, Bellows Falls, VT) red emitter, and an FF01-525/45-25 (Semrock) green emitter. Laser-scanning Dodt contrast was achieved by collecting the laser light after the spatial filter with an amplified diode (PDA100A-EC, Thorlabs). Our in-house jScan software (https://github.com/pj-sjostrom/jScan; https://zenodo.org/doi/10.5281/zenodo.10430989) running in Igor Pro 8 or 9 was used to create 2P optogenetic stimulation patterns and to acquire 2P imaging data, via PCI-6110 boards (NI, Austin, TX, USA).

### 2P Optomapping

To identify candidate presynaptic neurons in the FOV, a 2P image of mRuby fluorescence (512×512 pixels at 1.93 pixels/µm, average of 2 frames) was slowly acquired in jScan at 8 ms per line to ensure that scanner voltage command and scanner pointing location matched. The scanner position feedback signal was thus not used.

Fluorescent cell bodies in the image were automatically detected in jScan, with the occasional software identification mistake adjusted by the user. These coordinates were used to automatically place Archimedean spiral trajectories (15 µm diameter, ∼1 µm revolution spacing) over each cell body. For each cell location, spirals were next converted from pixel coordinates back to scanner voltage commands (spiral arc step 15 mV and separation 20 mV in command voltage terms). Each candidate presynaptic cell received a burst of three spirals, 7 ms duration each, at 30 Hz, with the 2P laser gated by the GVS011/M galvanometric mirror shutter. These spiral scan bursts on individual cells were separated by 200 – 500 ms, with cell-to-cell fly time set to 5 ms. Candidate presynaptic cells in an FOV were thus sequentially stimulated twenty times every 30-40 seconds while the postsynaptic cell was whole-cell recorded in current-clamp mode. At the end of each stimulation sweep, five 1.3-nA depolarizing pulses at 50 Hz followed by a 250-ms-long 25-pA hyperpolarizing pulse were delivered to the patched cell to monitor membrane potential, input resistance, and its continued ability to spike. Recordings with more than 30% change in input resistance or more than 8 mV change in resting membrane potential were deemed to be unstable and discarded. Electrophysiology and optogenetic stimulation were synchronized by triggering. To create connectivity maps without gaps, FOVs overlapped slightly but were offset in z to avoid photostimulating the same candidate presynaptic cells more than once.

To semi-automatically detect EPSPs from candidate presynaptic cells, the twenty acquired sweeps of each FOV were combined. A Student’s t-test comparison of the average membrane potential during a 5-ms-long baseline period before spiral-scan onset and a 5-ms-long window 7-10 ms after the first of three spiral-scans was used to identify synaptic responses at the p < 0.05 significance level. Erroneous false positives and false negatives were user-corrected by visual inspection of the responses to the remaining two in the 30-Hz burst of three spiral-scans. For cells that were thus deemed connected, the amplitudes of EPSP_1,_ EPSP_2,_ and EPSP_3 i_n the 30-Hz train were determined by peak-searching the average of the 20 sweeps. Temporal summation was accounted for by valley-searching between the EPSP peaks. The CV of synaptic responses was determined from the standard deviation calculated from the 5-ms-long windows of the 20 sweeps, divided by the mean amplitude obtained by peak-searching the average trace.

In rare cases (<7% of connections) for which the first spiral-scan failed to elicit an EPSP but the subsequent two did, EPSP_2 w_as automatically considered EPSP_1,_ and EPSP_3 w_as taken as EPSP_2._ In other words, we assumed that in this scenario, the failure to elicit a response on the first spiral-scan was because the presynaptic cell was not optogenetically driven to spike, but temporal summation led to presynaptic spiking after the second and third spiral-scan.

Paired-pulse ratio (PPR) was calculated as the ratio of EPSP_2 t_o EPSP_1._ If the first spiral-scan failed to elicit an EPSP but the subsequent two did, PPR was calculated from the subsequent two pulses. To ensure robustness, PPR data points were omitted when the leading EPSP was smaller than 0.1 mV, similar to in our prior studies.^78^ However, these inputs still counted towards connectivity statistics and their amplitudes still contributed to statistical comparisons and density maps.

### Accounting for optogenetic stimulation artifacts

To validate the single-cell resolution of 2P spiral-scanning, we estimated the median intercell distance, *r̃*, of mRuby-positive neurons in n = 25 fields of view (FOV) sized 265×265 µm and found that *r̃* averaged 21 ± 0.4 µm. The density of neurons in mouse cortex has previously been estimated to *ρ* = 9.2×10^4^/mm^3^,^8^ which yields a mean inter-particle radius of 〈*r*〉∼ 1⁄*ρ*^1/3^= 22 µm. Since our z-axis resolution *w*_½_ was less than half compared to either of the radii *r̃* or 〈*r*〉, we concluded that we achieved single-cell resolution with an appreciable margin.^43^

We were concerned that optomapping would be sensitive to artifacts related to the acute slice preparation. Too close to the slice surface, connections would have been physically severed during dissection. Too deep into light-scattering brain tissue, 2P excitation might be insufficient, leading to loss of connections due to failure of presynaptic suprathreshold excitation. We thus expected that there might be an ideal depth range. To delineate this range, we optomapped the same FOV at several different z-axis depths (Figures S2A-S2D). We found that connectivity was indeed lower within 50 µm of the slice surface, as expected from dissection artifacts (Figure S2E). However, connectivity dropped off relatively weakly with increased depths (Figures S2E and S2F), suggesting that loss of 2P excitation deep into brain slices was a minor problem. We therefore targeted our optomapping experiments relatively deep, typically 97 ± 3 µm from the slice surface, based on a random sample of n = 25 cells (Figure S2F).

When the patched cell also expressed the opsin — which was the case for most PCs, but never for BCs or MCs — it was occasionally possible to inadvertently directly depolarize the recorded PC weakly while targeting nearby candidate presynaptic cells. To assess the severity of this potential problem and to find candidate solutions, we relied on pharmacological synaptic blockade to differentiate direct optogenetic depolarization from EPSPs (Figure S3A).

As expected, EPSPs were abolished by synaptic blockade, whereas direct optogenetic depolarization responses persisted (Figure S3A). Direct light-induced depolarization was instantaneous, whereas genuine EPSPs were naturally briefly delayed due to synaptic transmission as well as to presynaptic integration of opsin current (Figure S3B). Therefore, direct optogenetic stimulation could be differentiated from EPSPs based on the latency of the depolarization onset. If statistically significant depolarization in a 1-ms-long window after spiral-scan onset occurred across the 20 sweeps at the p < 0.05 level, it was therefore deemed to be the direct result of light stimulation (Figure S3B). As expected, direct stimulation responses had very small CV compared to genuine EPSPs as well as no appreciable short-term depression (Figure S3C and D). Direct stimulation furthermore only occurred for candidate presynaptic cells within ∼60 µm of the patched cell (Figure S3E-H). We could thus rely on a combination of recorded cell type (PC vs. BC/MC), depolarization onset latency, relative stimulation location, and the analysis of CV and short-term plasticity to determine whether stimulation at a given presynaptic cell location elicited only an EPSP, only direct depolarization, or both (Figure S3I).

In the ‘both’ scenario, we devised an approach to estimate the synaptic component. Under synaptic blockade, we found that the peak amplitude of direct depolarization was highly correlated with the baseline amplitude 1 ms after spiral scan onset (Figure S3J). To estimate connection strengths in the ‘both’ scenario, we used the baseline amplitude to predict the direct depolarization amplitude. The predicted direct depolarization amplitude was subtracted from the measured EPSP amplitude to estimate the connection strength. All connections that involved a direct depolarization element were also manually inspected.

### Peak Depolarization Latency

Since we observed short-term facilitation and temporal summation in MC EPSPs (Figure 7), we suspected that measuring the amplitude of the first response, i.e., EPSP_1,_ in a train of three underestimated excitation of MCs. An alternative approach would be to measure the peak depolarization, ΔV_peak,_ due to the temporal summation of the three EPSPs.

To measure ΔV_peak,_ we averaged the 20 sweeps and searched within a 120-ms time window from the spiral-scan onset for the maximum depolarization relative to the 5-ms-long baseline before spiral-scan onset. With the ΔV_peak m_etric, MC synaptic input density maps (Figure S8A) were spatially indistinguishably structured from their EPSP_1 c_ounterparts (Figures 3A, 4A, 5A). However, the excitation of MCs was as expected now markedly stronger (Figure S8B, C). We also compared the latency of peak depolarization in PCs, BCs, and MCs (Figure S8D).

### Connectivity Mapping

For each FOV, EPSPs were matched to their respective presynaptic cell (Figure 1E). FOVs were next arranged according to their microscope stage coordinates to create a connectivity map spanning the cortical thickness (Figure 1F). Neocortical layer boundary lines were drawn (see below). Connectivity maps were rotated to their straight-up position and an arbitrary 200-µm-wide vertical column was centered on the postsynaptic neuron to standardize connectivity comparisons across cells (Figure 1G). To generate individual connectivity density maps, cell-sized 2D Gaussians (sigma 10 µm) with peak amplitude scaled according to EPSP magnitude were summed up (Figure 1H).

Ensemble connectivity density maps (Figures 3A, 4A, 5A) were generated by averaging individual connectivity density maps for specific postsynaptic cell classes. When averaging individual maps, the neocortical layers naturally varied in thickness from recording to recording. To account for this variability, the average layer thickness for L2/3, L4, L5, and L6 were calculated, and then the layers of individual maps were standardized by slightly stretching or shrinking to account for the variable layer thickness, to permit averaging with matching cortical layer boundaries. This is why layer boundaries in ensemble heat maps have no error bars, yet layer boundaries vary slightly from one ensemble heat map to the next (e.g., Figures 3A, 4A, 5A)

Statistics were carried out across postsynaptic cell types, accounting for individual postsynaptic neurons nested in individual animals as random effects (see Statistics section). Data on connectivity, EPSP amplitude, and short-term synaptic dynamics were collected based on the presynaptic cell’s layer location (Figures 3B-C, 4B-C, 5B-C) or radial distance relative to the recorded postsynaptic neuron (Figure S7). Radial connectivity was averaged across postsynaptic cells for each binned radius and fitted with exponentials (Figure S7). Due to poorly converged exponential fits, one *τ* value was removed from the L2/3 and L6 PC categories each.

### Identification of Neocortical Layers

Neocortical layer boundaries were identified by mRuby fluorescence and Dodt contrast channels. Like before,^46^ we defined L5 by the presence of conspicuous PCs with large somata and thick apical dendrites, whereas PCs in L6 had thin apical dendrites and rounded somata. L4 was characteristically slightly darker and granular. L1 and white matter were relatively devoid of cells. L2/3 was defined by the presence of relatively small PCs. Layer boundaries were additionally informed by the Allen Mouse Brain Atlas^47^ and the Paxinos Mouse Brain Atlas.^48^

### Neuronal Morphometry

After each whole-cell recording, 2P image stacks were acquired (512×512-pixel slices at 1.9 pixels/µm, averaged over 2 frames, 8 ms/line, with slices separated by 2-3 µm). Excitation wavelength was 780 nm or 820 nm for Alexa 488 and Alexa 594, respectively. In addition to the biocytin histology and confocal imaging procedure described below, this 2P stack provided a complementary morphological cell identification method.

After acquiring the 2P image stack, the patch pipette was slowly removed, and the acute brain slice was fixed overnight in 4% paraformaldehyde (Millipore Sigma, St. Louis, MO) at 4°C. Sections were then stored in the dark, in 0.01 M phosphate buffered saline (Millipore Sigma), at 4°C for up to 3 weeks before staining.

Sections were washed 4 times for 5-10 minutes in 0.01 M Tris-buffered Saline (TBS) solution with 0.3% Triton-X, followed by a 1-hour incubation in a block solution composed of 0.01 M TBS, 0.3% Triton-X, and 10% normal donkey serum (NDS; 017-000-121, Jackson ImmunoResearch, West Grove, PA). Sections were subsequently incubated in immunowash solution (0.01 M TBS, 0.3% Triton-X, 1% NDS) supplemented with 1:200 Alexa Fluor 647- or Alexa Fluro 488-conjugated Streptavidin (Thermo Fisher) overnight at 4°C. The following day, sections were washed 4 times for 5-10 minutes in 0.01 M TBS, mounted on glass slides, and coverslipped using 40 µL ProLong Gold Antifade Mountant (Thermo Fisher).

Sections were imaged using a Zeiss LSM780 confocal laser-scanning microscope and Zen Software (Zeiss, Oberkochen, Germany) no later than 1 week after staining. Images were acquired using the Plan-Neofluar 20×/0.4 LD objective (Zeiss). A 633 nm He-Ne laser and an Argon Multi-line laser tuned to 488 nm were used to excite Alexa Fluor 647 and 488, respectively. 3D image stacks (708.5×708.5-pixel slices at 1.445 pixels/µm separated by 1.07 µm) were acquired and used for morphological reconstructions.

Confocal image stacks were contrast adjusted and converted to 16 bits in Fiji^98^ and then imported into Neuromantic^99^ for manual 3D tracing. Neocortical layers and image stack outlines were likewise manually labeled in Neuromantic.

Like we described before,^46,100^ quantification and statistical analysis of 3D reconstructed neurons were carried out using qMorph in-house software (https://github.com/pj-sjostrom/qMorph, https://doi.org/10.5281/zenodo.7853963) running in Igor Pro. Morphological reconstructions were rotated around the soma so that the pial surface was aligned in the “straight up” position (Figure S5). Maximum ascending and descending neurite distances were measured from the center of the soma (Figure S4C). To create density maps (Figure S5B), 2D Gaussians with sigma 25 µm aligned on compartment XY centers and with amplitude proportional to compartment size were summed to form a smoothed morphology 2D projection. To permit averaging across reconstructions, individual density maps were peak normalized. Axons and dendrites were density-mapped separately, normalized, and merged by logical OR. Color maps in Figure S5B are thus in arbitrary units but comparable across cell and compartment type. To create symmetrical density maps, reconstructions were mirrored, but all metrics were carried out on non-mirrored data.

Convex hulls of individual reconstructions were constructed from 2D projections of axonal and dendritic arbors, using a Jarvis walk. A convex hull of all convex hulls of a given cell type was used to generate an ensemble convex hull (Figure S5B). Sholl analysis^101^ was carried out by aligning reconstructions on somata, converting to radial coordinates, moving outwards from somata in 6.5-µm steps, and counting the number of compartments crossing a given radius (Figure S5C). Sholl diagrams were averaged without normalization.

### Cell Classification

For each optomapped cell, we assessed spike patterns from current-clamp input/output curves using 500-ms-long current pulses starting at -10 pA and ending at 200-600 pA, in 10-20-pA steps (Figure S5E). Spike width, height, and threshold were measured from the smallest current injection trace that elicited spikes, i.e., the rheobase trace (Figure S4C and S5F).

Instantaneous frequency was measured from the first two spikes in the rheobase trace. Accommodation was calculated as the ratio of instantaneous frequency for the first and last spike pairs in the rheobase spike train. Metrics such as frequency, accommodation, etc., that required more than one spike, were measured from the rheobase +20 pA trace if too few spikes were elicited in the rheobase trace. These metrics are summarized in Figure S4C.

Using JMP Version 16 (SAS Institute Inc., Cary, NC), principal component analysis was carried out for the electrophysiological and morphological parameters highlighted in Figure S4C (Figure S4B). The first two principal components were subsequently used for agglomerative hierarchical clustering using Ward’s minimum-variance method (Figure S4A). Due to optical sectioning, maximum ascending and descending neurite measurements were missing for a subset of cells (19 of 38 PCs; 2 of 21 BCs), so missing values were imputed for clustering purposes.

However, reported averages (Figures S4C, S5C and D) do not include imputed values. The number of clusters was automatically determined by JMP’s default approach: detecting a sharp change in linkage as a function of branch number. This approach revealed three distinct classes in L2/3, L5, and L6 (Figure S4A) that corresponded well to PCs, BCs, and MCs.^18^

### Comparison with Paired Patch

For validation, we compared L5 optomapping within a 100 µm radius of the postsynaptic cell (Figure 5A, dashed circle) to L5 paired-recording data that we previously acquired.^46^ In that study, the paired-recording data set was acquired in acute V1 slices obtained from P11-P20 mice. PC→PC and PC→BC data was obtained from C57BL/6 mice as well as from the G42 mouse line,^102^ i.e., the same genetic background as Emx1-Cre mice^38^ used here. However, the PC→MC data was acquired by targeting fluorescent cells in the GIN mouse line, which were bred on the original albino FVB/N background.^103^ Since albinism leads to visual system miswiring in mice and other species,^104^ this is a potential caveat.

Whereas optomapping can only find unidirectional connections, bidirectional connections contributed to the PC→PC paired-recording connectivity data set.^26^ To enable comparison with optomapping, the PC→PC paired-recording connectivity, which was measured to 162/1208 = 13.4%, was therefore adjusted to 81/682 = ∼11.9% by considering only one direction for each measured paired recording. Thus, the number of unidirectional connections were taken as half of the measured unidirectional connections plus half of the measured bidirectional connections, 2*N*(1 − *p*)*p*/2 + *Np*^%^/2 = 81, while the number of unconnected cells were taken as half of the measured unconnected pairs plus half of the measured unidirectional pairs, *N*(1 − *p*)^%^/2 + 2*N*(1 − *p*)*p*/2 = 601, where *N* is the total number of tested connections and *p* was implicit from the number of measured unconnected pairs, *N*(1 − *p*)^%^ = 1208. Half of the measured unconnected pairs were thus considered never tested. However, to compare EPSP amplitudes, the entire PC→PC paired-recording connectivity data set was used. The PC→BC and PC→MC paired-recording data sets were not adjusted, since these were by default only considered in one direction, much like optomapping.

### Statistics

Results are reported as mean ± standard error of the mean (S.E.M.), unless stated otherwise (Figure 8). Statistical tests were conducted in Igor Pro 9, unless otherwise stated. We observed no significant difference in connectivity and EPSP amplitude between the sexes (connectivity: males, 8.12% ± 0.30%; females, 7.34% ± 0.31%; p = 0.35, GLMM; amplitude: males, 0.67 ± 0.04 mV; females, 0.84 ± 0.07 mV; p = 0.88, LMM). Therefore, data from both sexes were pooled.

To determine the best-fit statistical model to EPSP amplitude distributions (Figure 2C and S6), we conducted AIC model selection in JMP 17, which compared the fit of 11 continuous distributions (normal, Cauchy, Student’s *t*, sinh-arcsinh, exponential, gamma, log normal, Weibull, normal 2 mixture, normal 3 mixture, and Johnson’s *S_U_*_)_. For each cell type, we reported the AICc weights, which represent the relative likelihood of a model, for a log-normal distribution in Figure S6B. In JMP 17, we used the Anderson-Darling test to determine how well the log-normal distribution describes the EPSP amplitudes of inputs onto each cell type^105,106^. The test statistics, *A^2^*, and corresponding p values are reported in Figure S6B.

Synaptic connectivity rates, EPSP amplitudes, and PPRs were compared between cell types and input layers using GLMMs for connectivity rates or LMMs for EPSP amplitudes and PPRs (Figures 3B, 3C, 4B, 4C, 5B, 5C, and 7E), running in RStudio 2023.06.1.524 (Posit Software, Boston, MA).^107^ We used mixed models because individual data points (i.e., individual connections) recorded from a single postsynaptic neuron or from the same animal were not independent of each other. Therefore, individual postsynaptic neurons nested in individual animals were included as random effects. To assess connectivity rates, all tested connections were labeled as either unconnected or connected, resulting in a binomial error structure. GLMM statistics were estimated using the mixed() function from the afex package,^108^ which calls on the glmer() function from the lme4 package.^109^ An F test was carried out to test the significance of fixed effects (presynaptic layer location and cell type, or sex) using the fitted model as the argument in the anova() function from the stats package.^110,111^ If fixed effects were significant, pairwise comparisons were carried out using the emmeans() function from the emmeans package,^112^ which uses the Tukey method to adjust the p-values for multiple comparisons. Since EPSP amplitudes and PPRs were found to be log-normally distributed, we fit LMMs, which requires normally distributed data, on logged EPSP amplitudes or logged PPRs using the lme() function from the nlme package.^110^ The lme() function uses the restricted maximum likelihood method to estimate LMM statistics. As described above, an F test was carried out using the anova() function and if fixed effects were significant, pairwise comparisons were carried out using the emmeans() function.

Connectivity radial τ was compared between the 3 cell types in L2/3, L5, and L6 using the Kruskal-Wallis *H* test (Figure S7). We also pooled radial τ from BCs and MCs to compare against PCs. The Wilcoxon-Mann-Whitney two-sample rank test was used for two-sample comparisons, with Zar’s normal approximation if the combined n’s were larger than 40. Optomapping validation with paired-patch connectivity^26^ was carried out with chi-square tests in Excel.

The statistical significance of high-order connectivity patterns (Figure 8) was determined using a Monte-Carlo approach. For L6 PCs, all presynaptic cells tested in a ROI (within the column and within L6) were shuffled independently for the two postsynaptic neurons and the number of shared inputs were counted across 10,000 runs. The resulting distribution of shared-input counts was used to determine the probability of the experimentally encountered shared-input count. For postsynaptic cells in L2/3 (Figure 7), the ROI was restricted to inputs within the column and above the L4/L5 boundary, like in the prior literature.^65^ By within column, we mean the intersection of 200-µm-wide columns centered on the two postsynaptic cells, although the union of the two columns gave similar results. Although we report results for 10,000 runs, we note that 100,000 runs gave indistinguishable significance levels.

To determine the magnitude of over or under representation of shared inputs, we relied on Monte-Carlo bootstrap.^113^ We randomly subsampled 60% of all inputs in the ROI at hand and shuffled them in one of two ways. In one scenario, inputs were randomized the same for the two postsynaptic cells, thus preserving the structural overlap of inputs as well as the connectivity rates while randomizing connectivity onto individual neurons. In the other scenario, inputs were randomized independently for the two postsynaptic cells, thus disrupting both structural overlap of inputs as well as connectivity onto individual cells, while preserving connectivity rates. As a measure of surplus or deficit in overlap, the distribution from the first scenario was ratioed to that of the second, with standard deviations scaled accordingly. We pooled distributions across postsynaptic pairs. We report results for 10,000 runs for each scenario but we found no appreciable difference when using 100,000 runs. We tried subsample sizes ranging from 50% to 80% but found no differences.

## KEY RESOURCES TABLE

| REAGENT or RESOURCE | SOURCE | IDENTIFIER |
| --- | --- | --- |
| **Antibodies** |  |  |
| Alexa Fluor 488-conjugated streptavidin | ThermoFisher Scientific | S32354 |
| Alexa Fluor 647-conjugated streptavidin | ThermoFisher Scientific | S32357 |
| **Bacterial and virus strains** |  |  |
| AAV9-CAG-DIO-ChroME-ST-P2A-H2B-mRuby3 | Addgene | 108912 |
| AAV9-mDlx-GFP-Fishell-1 | Addgene | 83900 |
| **Chemicals, Peptides, and Recombinant Proteins** |  |  |
| D/L-AP5 | Hello Bio | A5282 |
| NBQX | Hello Bio | S5567 |
| Alexa Fluor 488 hydrazide | ThermoFisher Scientific | F14221 |
| Alexa Fluor 594 hydrazide | ThermoFisher Scientific | A10438 |
| **Experimental Models: Organisms/Strains** |  |  |
| Emx1<sup>IRES</sup>Cre/IRES Cre | The Jackson Laboratory | RRID:IMSR_JAX:005628 |
| **Software and Algorithms** |  |  |
| Igor Pro | https://www.wavemetrics.com/ | Igor Pro 9 |
| FIJI | https://imagej.net/software/fiji/ | V2.9.0 |
| Neuromantic | https://www.reading.ac.uk/neuromantic/body_index.php | V1.7.5 |
| JMP | https://www.jmp.com/en_is/home.html | V 17 |
| RStudio | https://posit.co/download/rstudio-desktop/ | 2023.06.1.524 |

---

**Figure S1.**
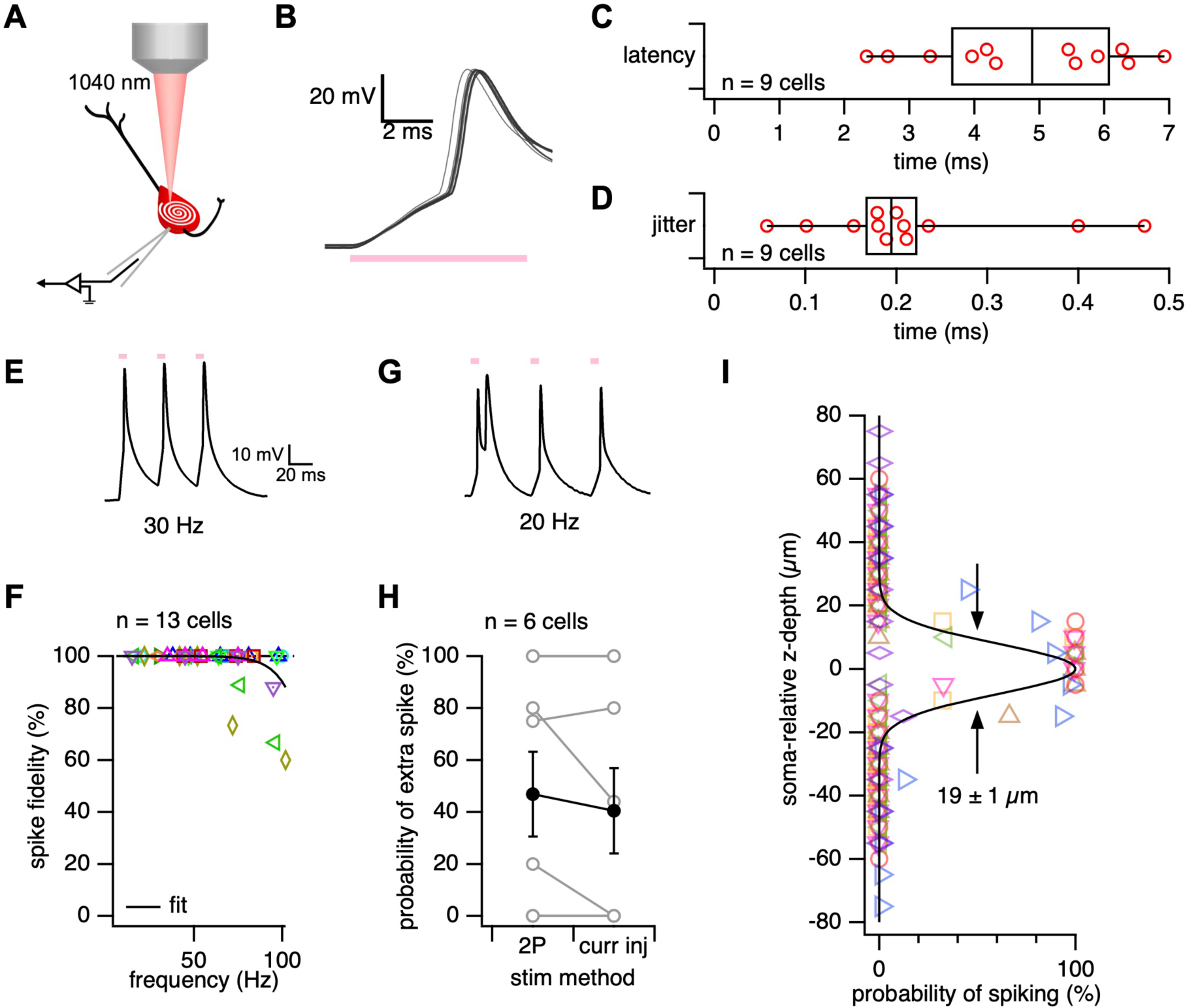
2P spiral scanning reliably drives spikes in ChroME-expressing neurons with single-cell spatial and sub-millisecond temporal resolution. Related to Figure 1. (A) We whole-cell recorded from a ChroME-expressing PC while activating it with 1040-nm spiral-scans over the soma. (B, C, and D) Spiral-scans reliably induced APs within milliseconds and repeated spiral scans drove time-locked APs with sub millisecond jitter. Latency was measured as the time between spiral scan onset and AP inflection point. Jitter was measured as the difference between the minimum and maximum latency resulting from repeated spiral scans of the same neuron. The box plot denotes the median, first, and third quartile, and the whiskers represent the minimum and maximum of the ensemble. (E and F) Spiral scans can drive ChroME-expressing PCs to spike reliably at frequencies up to ∼70 Hz. For optomapping experiments, we drove neurons with a 30-Hz train of 3 spiral scans, as shown in (E). Different symbols in (F) represent individual PCs. (G and H) Spiral scanning occasionally induced double spikes, however the probability of spiral-scan-induced double spikes was indistinguishable from the probability of double spikes induced by 5-ms-long current pulses, which are typically used in paired recordings (2P spiral-scans, 46.9% ± 16%; current injection, 40.5% ± 16%; paired t-test, p=0.24). Black markers: Mean ± SEM. (I) Spiral scans induced APs with single-cell spatial resolution as PCs could not be reliably driven to spike if spiral scans were targeted more than ∼10 µm away from the soma.

**Figure S2.**
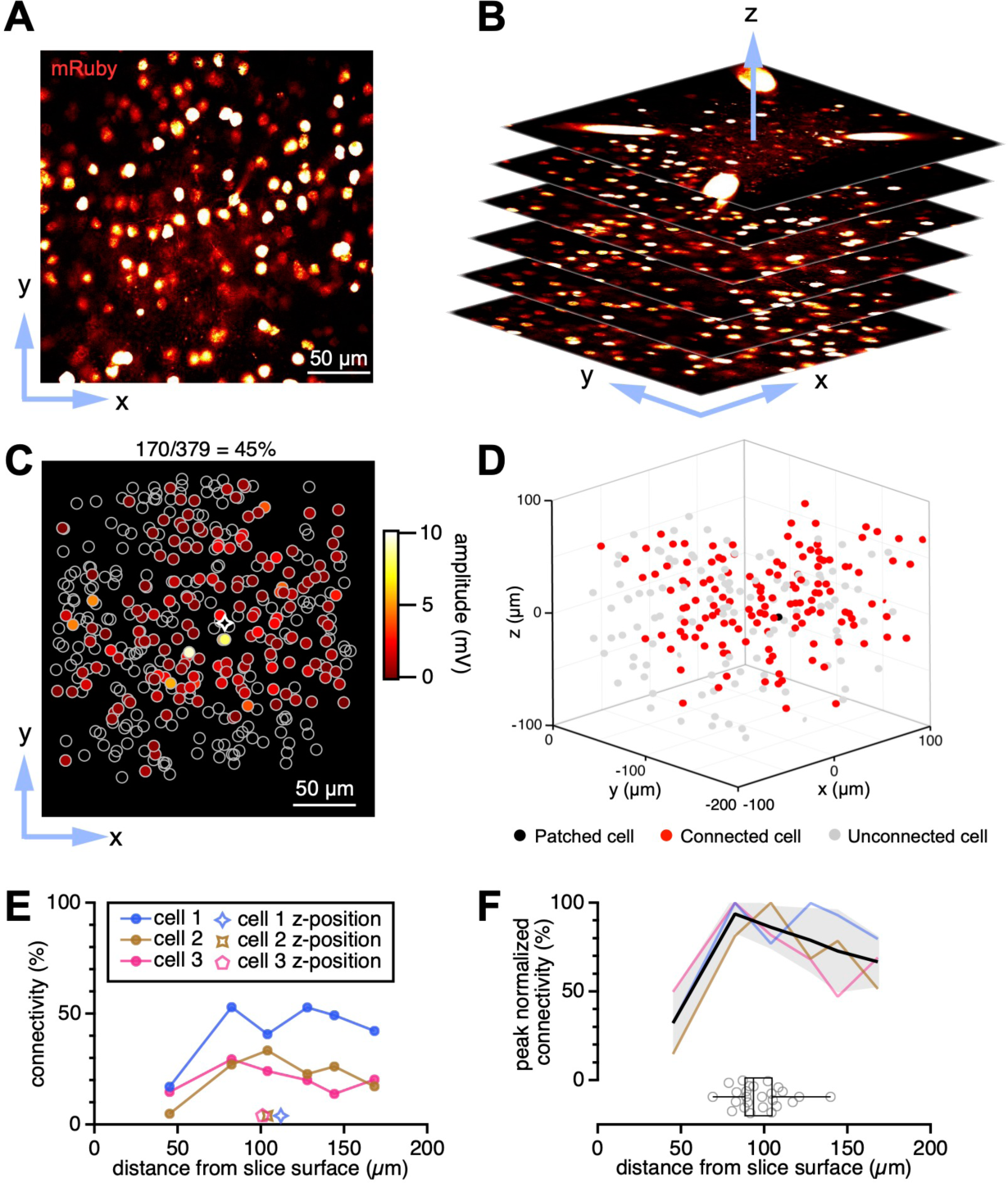
Reliable connectivity rates below 75 µm slice depth. Related to Figure 1. (A) mRuby3 fluorescence indicating cells which express ChroME in one microscope FOV. (B) Optomapping was carried out in a z-stack of microscope FOVs. (C) Flattened view of candidate presynaptic neurons (open circles) and connected excitatory inputs (colored circles) in the z-stack shown in (B). Connected neurons were color-coded by connection strength. White diamond: postsynaptic neuron. (D) Of the candidate presynaptic cells in (B) and (C) that were optomapped at different depths, some gave rise to EPSPs (red) whereas some did not (grey) in the patched postsynaptic neuron (black circle). (E) The connectivity rate of three postsynaptic cells (star symbols) appeared to increase with optomapping depth. Cell 1 (blue trace) is the example from (C) and (D). (F) Peak-normalized rates (colored lines) from (E) indicated that synaptic connections were reliably detected at slice depths of 75 µm or more. Inset box plot: The optomapping depth of 25 cells randomly selected from our dataset suggests our connectivity rates are reliable. Black line: Mean peak normalized connectivity rate. Grey shading: SEM peak normalized connectivity rate.

**Figure S3.**
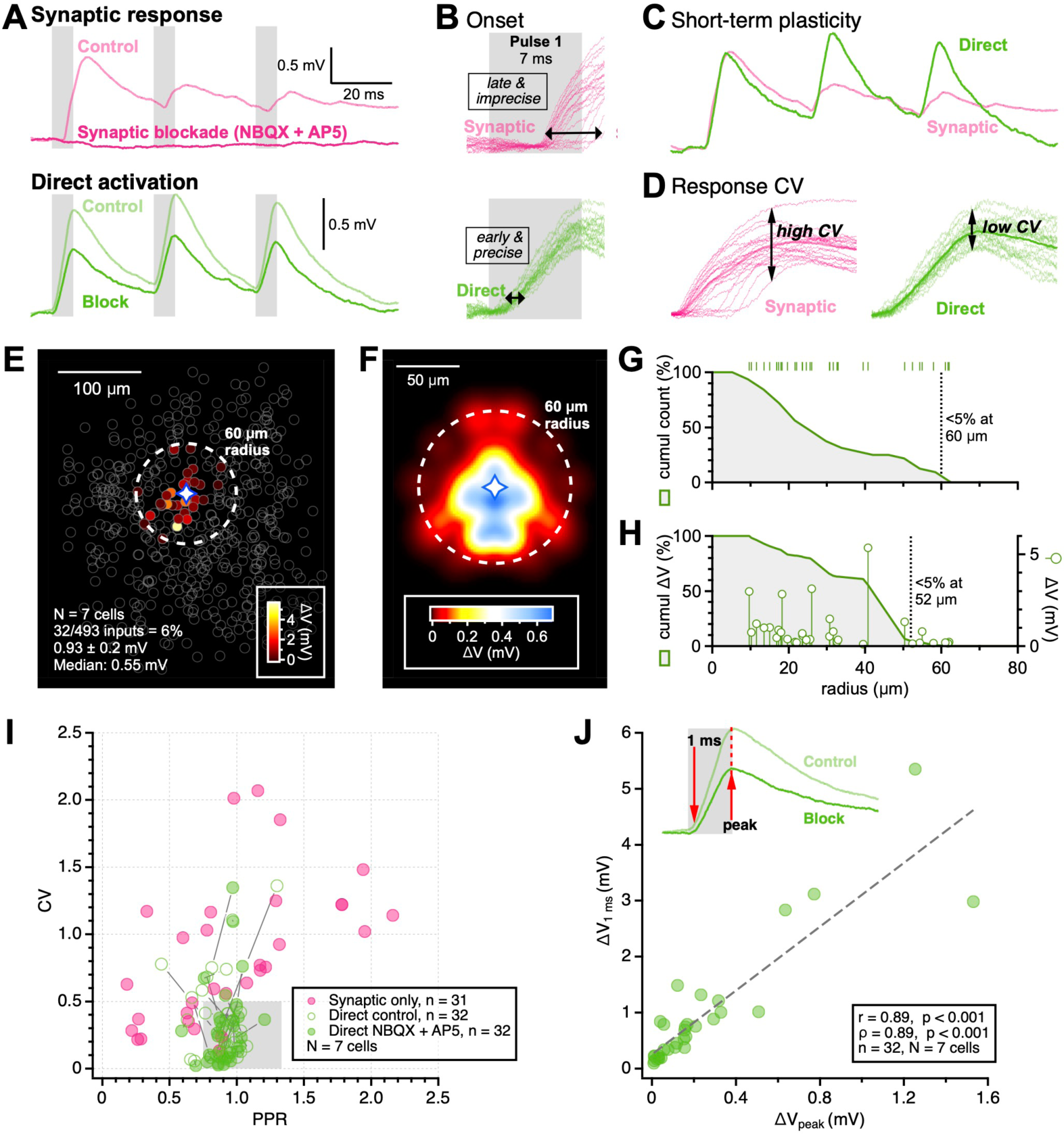
Differentiating optogenetic artifacts from EPSPs. Related to Figure 1. (A) Sample responses in a patched PC elicited by light-induced synaptic transmission, and light-induced direct depolarization. Light-induced EPSPs were completely abolished by NBQX and AP5 while light-induced direct depolarization was slightly reduced. (B) Light-induced EPSPs showed greater latency and jitter in response to spiral-scans than light-induced direct depolarization, where responses occurred instantaneously. (C) Light-induced EPSPs were generally short-term depressing, whereas light-induced direct depolarization showed no short-term plasticity. (D) Light-induced EPSPs showed more variable response amplitudes, or greater CV, than light-induced direct depolarization. (E) We visualized the location spiral-scans which induced direct responses in the recorded neuron by aligning the location of recorded PCs (diamond) and color-coded responses by amplitude. We found that direct light-induced responses only occurred when spiral-scanning candidate presynaptic cells within ∼60 µm of the patched PC (diamond). (F) The density map of spiral-scan induced direct responses showed that the spatial region around a neuron that was sensitive to direct light activation was roughly pyramidal shaped. We mirrored the location of responses across the vertical axis to generate a symmetrical density map. Diamond: patched PC. (G) Less than 5% of the instances of light-induced responses occurred outside a 60 µm radius of the patched cell soma. (H) The majority of depolarization elicited by direct light activation occurred within 52 µm of the patched cell soma. (I) Light-induced direct depolarization typically exhibited low CV and little short-term plasticity compared to light-induced EPSPs. The grey box denotes CV and PPR values which could predict responses that are purely due to light-induced direct depolarization with ∼80% accuracy. (J) In ACSF supplemented with NBQX and AP5, we studied the dynamics of light-induced depolarization. We could predict the peak amplitude of light-induced depolarizations based on the baseline amplitude at 1 ms after spiral-scan onset. Therefore, in cases where both light-induced depolarization and synaptic transmission occurred, we subtracted the computed amplitude of light-induced depolarization from the peak depolarization amplitude to estimate the EPSP amplitude (red dashed line).

**Figure S4.**
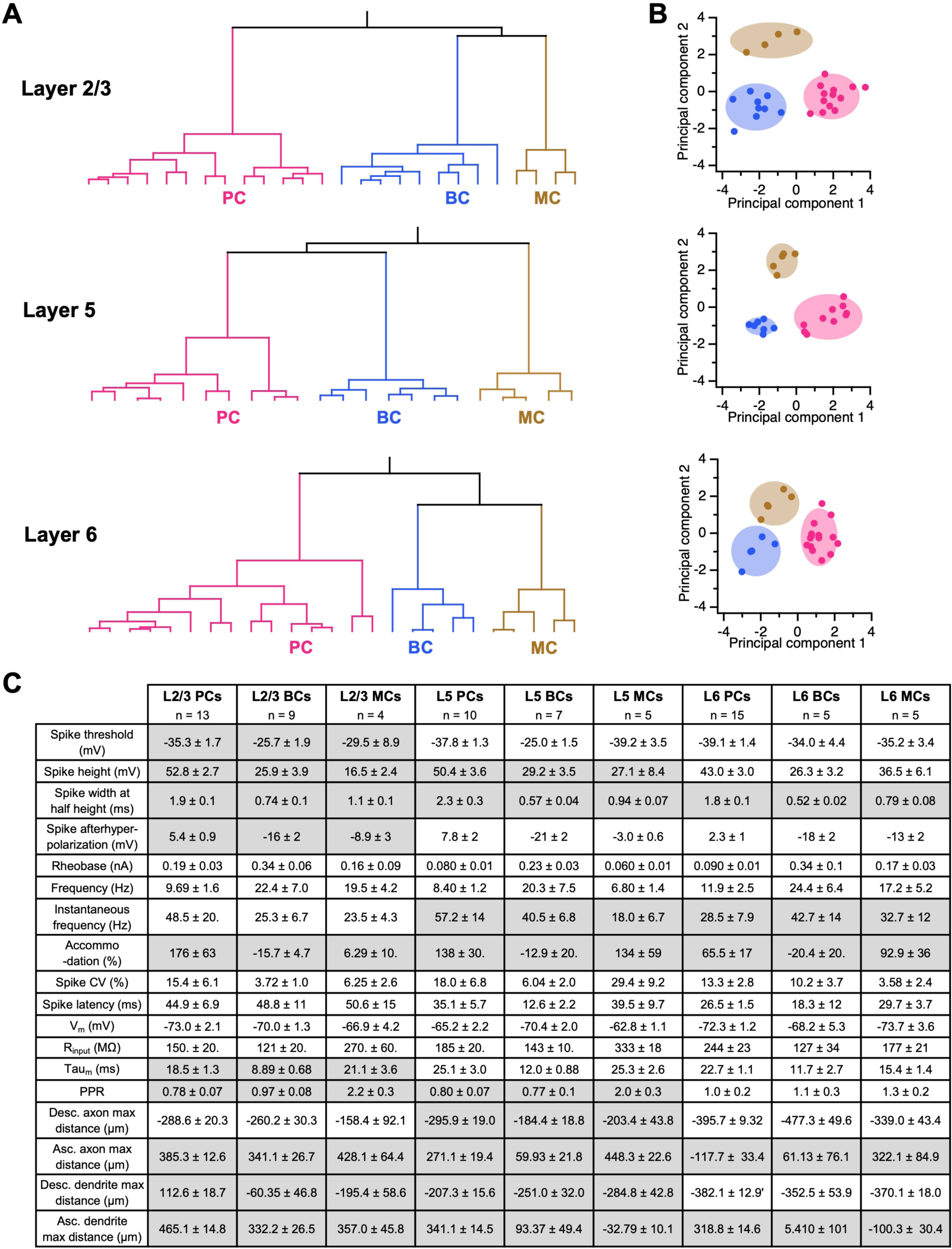
PC, BC, and MC clustering was robust. Related to Figures 2 – 7. (A) Patch-clamped cells were segregated by agglomerative hierarchical clustering on a combination of electrophysiological and morphological properties (see below). Clusters were automatically determined from a sharp change in linkage (transition from black to colored lines), which automatically segregated recorded cells from L2/3 (top), L5 (middle), and L6 (bottom) into three classes — PCs, BCs, and MCs. (B) The first two principal components of electrophysiological and morphological characteristics (see below) obtained from patch-clamped cells in L2/3 (top), L5 (middle), and L6 (bottom) indicated three distinct clusters that corresponded to PC, BC, and MC classes. Shaded areas: µ ± 2σ. (C) Electrophysiological and morphological properties of patch-clamped cells were distinct for different cell classes but relatively similar across layers for the same class. Numbers are Mean ± SEM. Grey boxes: parameters used for hierarchical clustering.

**Figure S5.**
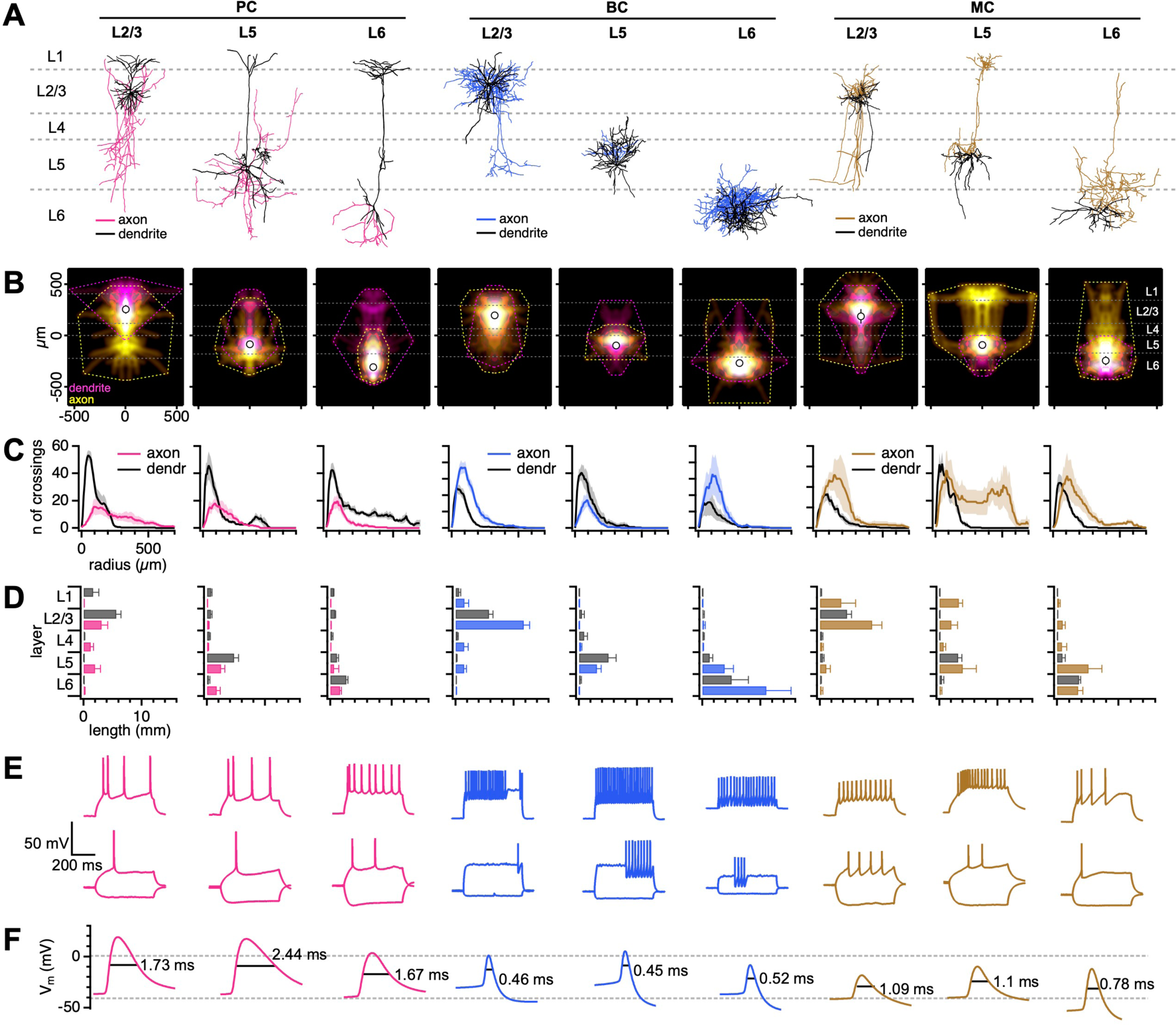
Distinct morpho-electrophysiology of PCs, BCs, and MCs. Related to Figures 2 – 7. (A-D) PCs, BCs, and MCs across L2/3, L5, and L6 exhibit distinct morphologies. PCs had a prominent ascending apical dendrite as well as a descending primary axon. The majority of BC neurites were restricted to the same layer as the cell soma. MCs had an ascending axon, which was most prominently seen in L5 and L6 MCs. Sample morphologies are shown in (A). Ensemble density maps for axonal and dendritic compartments are shown in (B). In each density map, convex hulls illustrate maximum extents of axonal (yellow dotted line) and dendritic (magenta dotted line) compartments. Open circles: average soma position. Horizontal white dotted lines: neocortical layer boundaries. Each ensemble Sholl diagram in (C) shows the number of axonal or dendritic branch crossings at a given radial distance from the soma. The total length of axonal and dendritic arborizations in each neocortical layer is shown in (D). (E) We observed distinct electrophysiological characteristics between PCs, BCs, and MCs. However, cells of the same type while cells of the same class in different layers exhibit similar electrophysiological characteristics. Sample electrophysiological responses evoked by a hyperpolarizing current step (middle), the response at rheobase (middle), the response evoked by a rheobase + 40 pA stimulus (top). (F) PCs, BCs, and MCs spikes were distinctly shaped. PC spikes were relatively wide and typically peaked above 0 mV; BC spikes were relatively narrow, with half-widths of ∼0.5 ms; MC spikes rarely peaked above 0 mV.

**Figure S6.**
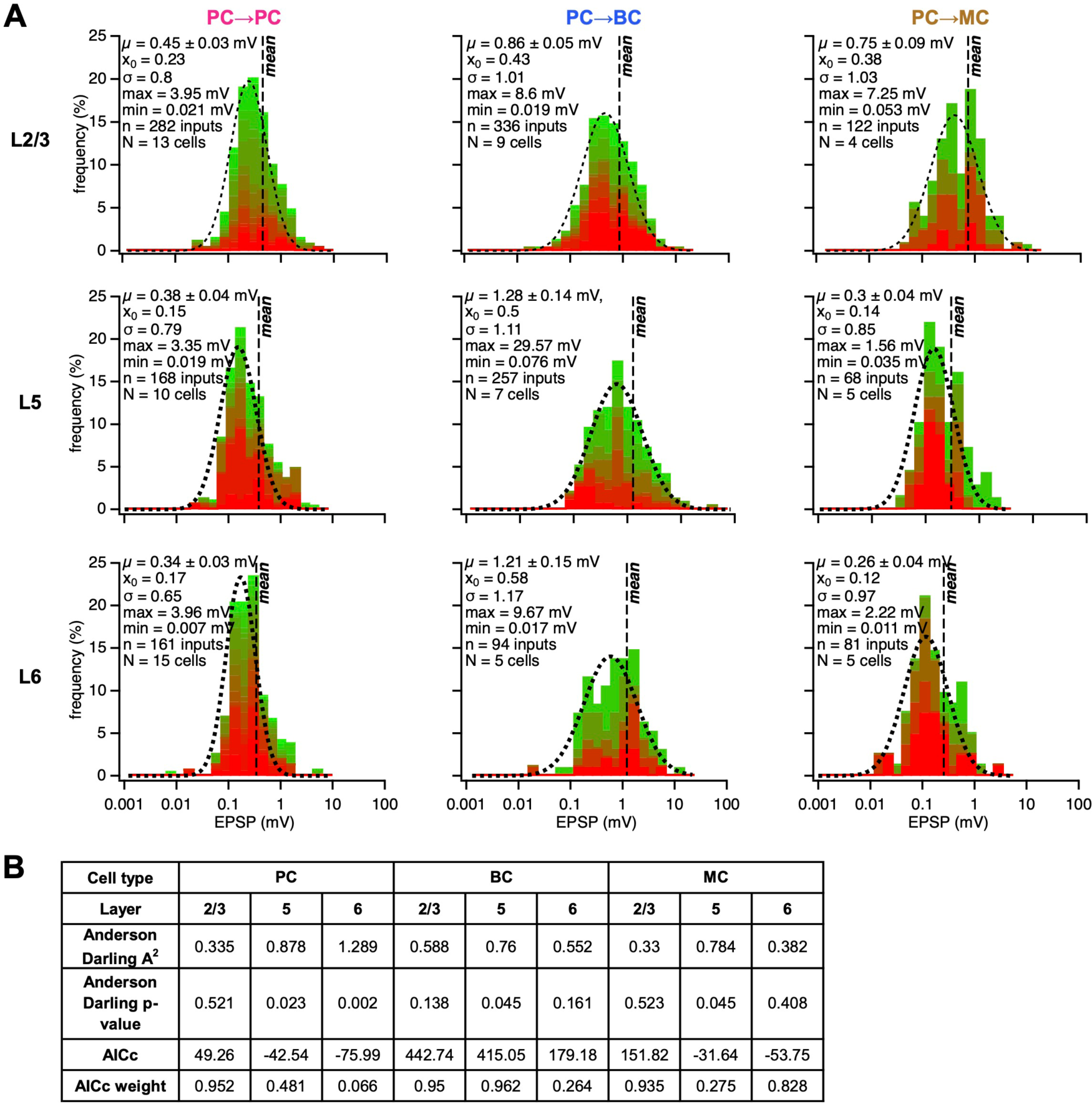
Excitatory inputs distributed log-normally. Related to Figure 2. (A) Ensemble EPSP amplitudes of PC, BC, and MC classes in L2/3, L5 and L6 distributed log-normally. Different colors represent individual cells. (B) When comparing 11 distribution types (see Methods), the Akaike information criterion (AICc) value and weight indicated that the log-normal distribution was the best fit for all EPSP amplitude datasets, even for groups with Anderson Darling p value < 0.05. In conclusion, the best statistical model of synaptic weight distributions was the log-normal.

**Figure S7.**
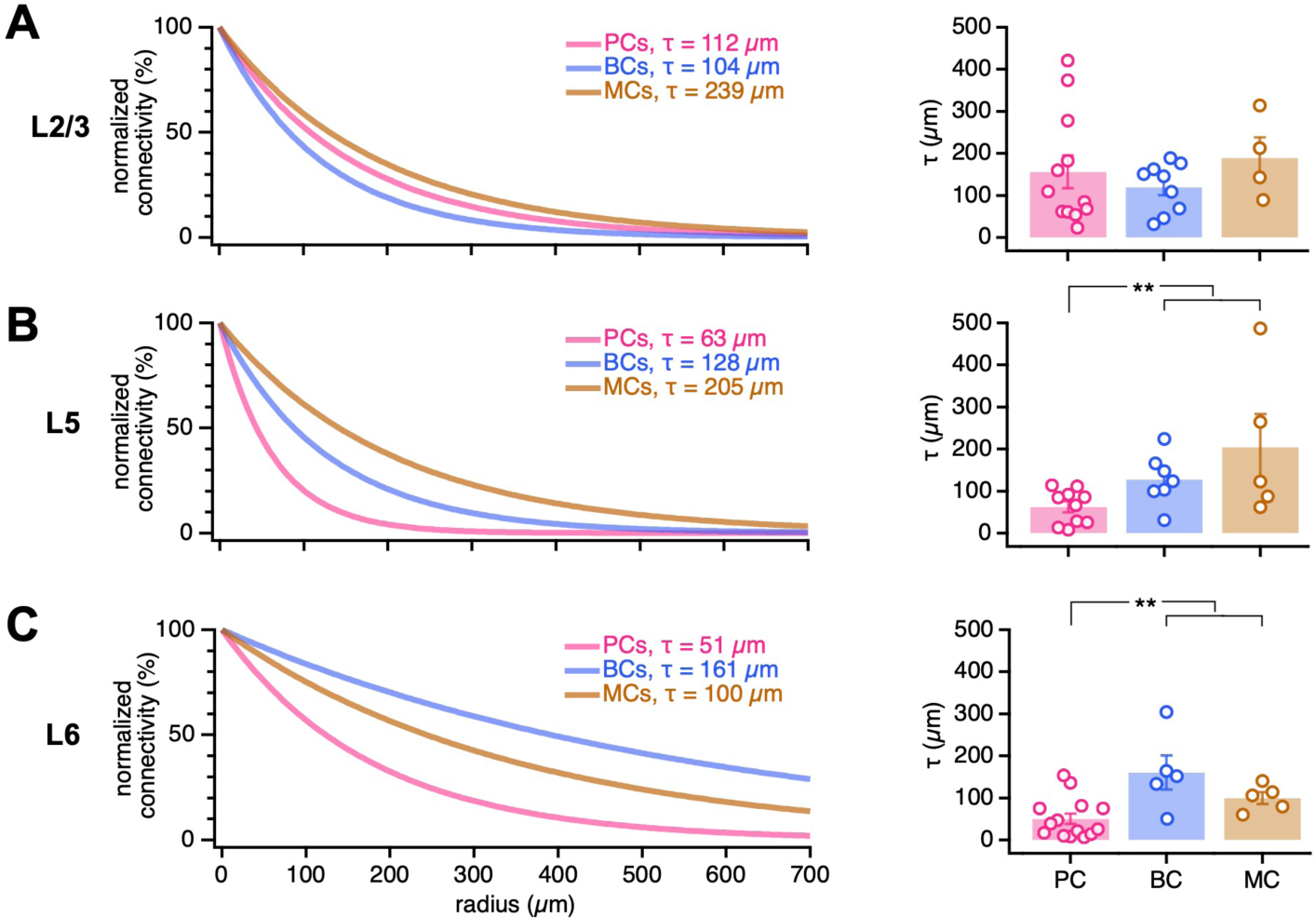
Subgranular interneurons received excitatory input from greater distances than PCs. Related to Figures 3 – 5. (A) To measure the distance-dependent decay of connectivity, *τ*, we relied on exponential fits to radially binned connectivity rates (see Methods). For L2/3, we found no differences across PCs, BCs, and MCs (p = 0.75, Kruskal-Wallis) even if we pooled BCs and MCs (pooled 141 ± 21 µm), like in mouse auditory cortex.^71^ (B, C) For L5 and L6, *τ* was greater in BCs and MCs than in PCs (L5 pooled interneurons, 160 ± 35 µm; p < 0.05, Kruskal-Wallis; L6 pooled interneurons, 130 ± 23 µm, p < 0.05, Kruskal-Wallis). This demonstrates that connectivity died off more slowly with distance for interneurons than for PCs, at least in subgranular layers. Mean ± SEM. \*\**P < 0.01*

**Figure S8.**
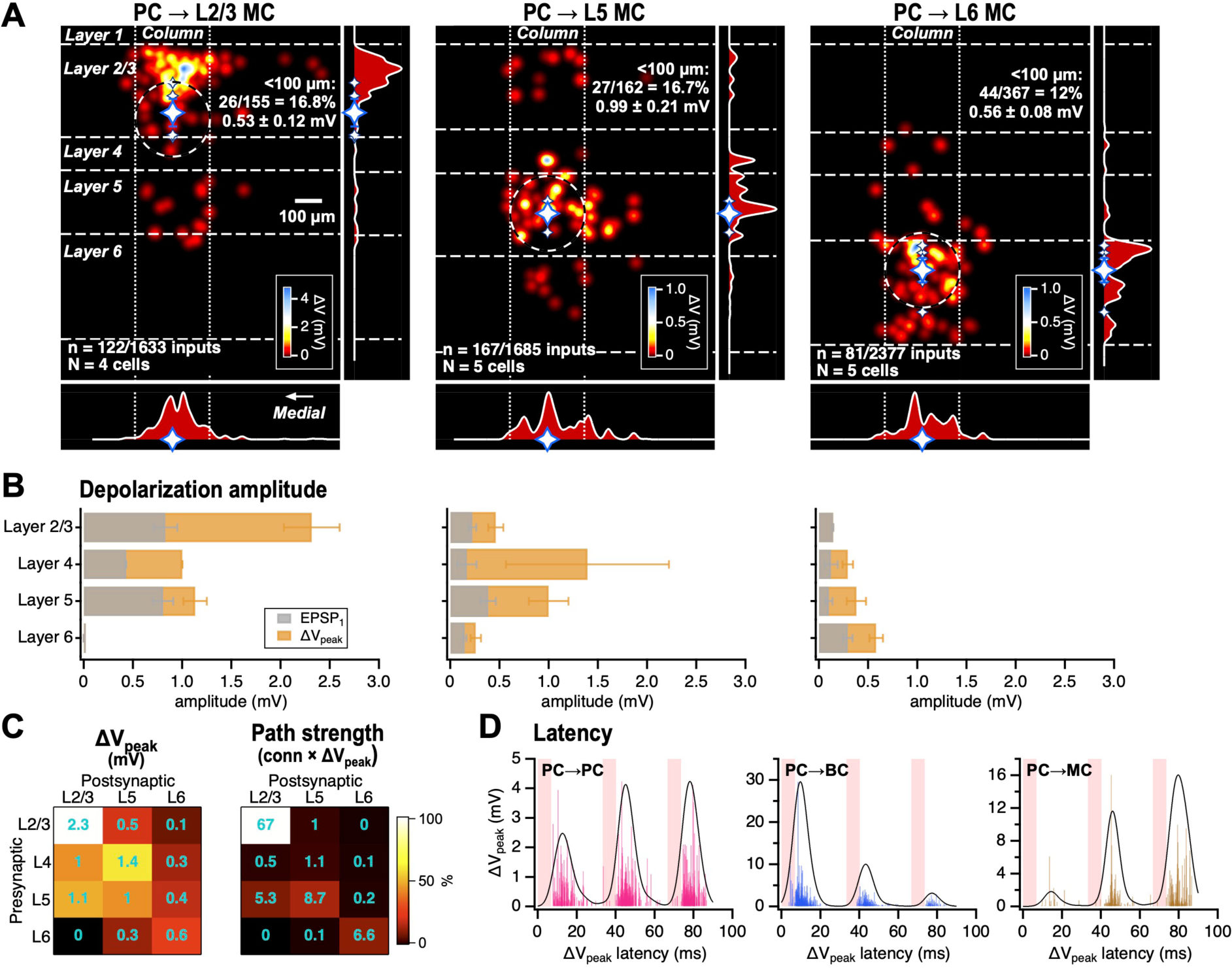
Peak excitation of MCs occurs with relatively long latency. Related to Figures 3 – 5. (A) To better measure the excitation of MCs, we relied on the peak depolarization, ΔV_peak,_ due to the temporal summation of the three EPSPs evoked at 30 Hz (Figure 1D). With this approach, synaptic input density maps for L2/3 (left), L5 (middle), and L6 MCs (right) were spatially structured indistinguishably from their EPSP_1 c_ounterparts (Figures 4A, 5A, 6A). Note, however, that the density map amplitudes were much larger with this metric. (B) The ΔV_peak m_etric highlighted a potent excitation of MCs. Grey bars show the EPSP_1 a_mplitude (as in Figures 4B, 5B, 6B), while the orange bars show ΔV_peak a_mplitude. Dynamically, the excitation of MCs is therefore appreciably stronger than that of PCs (Figures 4B, 5B, 6B). (C) When comparing PC→MC pathways across different input and output layers, using ΔV_peak s_hifted synaptic efficacy weights further towards the L2/3 PC→L2/3 MC path. While ΔV_peak r_evealed stronger overall excitation of MCs, the spatial structure of ΔV_peak p_ath strengths was similar to that of EPSP_1 (_Figure 7C), where within-layer communication dominated, and interlayer communication was largely biased towards L5→L2/3. When comparing across cell types, using ΔV_peak f_urther highlighted that the excitation of MCs — especially in the above pathways — is stronger than that of PCs (Figure 7B). Color code denotes within-matrix normalized values, which highlight differences within a target cell type. Cyan numbers indicate absolute values, which enables comparison across target cell type. (D) As expected,^21-23^ the structure of peak amplitude, ΔV_peak,_ versus its latency suggested early firing of BCs and late firing of MCs.^21-23^ Surprisingly, a relatively even temporal distribution of ΔV_peak e_merged for PCs across the three spiral scans (pink bars) delivered at 30 Hz (Figure 1D). Continuous lines are a peak-normalized sum-of-Gaussians, with individual Gaussians scaled by ΔV_peak a_nd with half-width 7 ms (same as spiral scan duration), to illustrate the average outcome for each target cell type. Mean ± SEM.

